# *In silico* approach toward the identification of unique peptides from viral protein infection: Application to COVID-19

**DOI:** 10.1101/2020.03.08.980383

**Authors:** Benjamin C. Orsburn, Conor Jenkins, Sierra M. Miller, Benjamin A Neely, Namandje N Bumpus

## Abstract

We describe a method for rapid in silico selection of diagnostic peptides from newly described viral pathogens and applied this approach to SARS-CoV-2/COVID-19. This approach is multi-tiered, beginning with compiling the theoretical protein sequences from genomic derived data. In the case of SARS-CoV-2 we begin with 496 peptides that would be produced by proteolytic digestion of the viral proteins. To eliminate peptides that would cause cross-reactivity and false positives we remove peptides from consideration that have sequence homology or similar chemical characteristics using a progressively larger database of background peptides. Using this pipeline, we can remove 47 peptides from consideration as diagnostic due to the presence of peptides derived from the human proteome. To address the complexity of the human microbiome, we describe a method to create a database of all proteins of relevant abundance in the saliva microbiome. By utilizing a protein-based approach to the microbiome we can more accurately identify peptides that will be problematic in COVID-19 studies which removes 12 peptides from consideration. To identify diagnostic peptides, another 7 peptides are flagged for removal following comparison to the proteome backgrounds of viral and bacterial pathogens of similar clinical presentation. By aligning the protein sequences of SARS-CoV-2 field isolates deposited to date we can identify peptides for removal due to their presence in highly variable regions that may lead to false negatives as the pathogen evolves. We provide maps of these regions and highlight 3 peptides that should be avoided as potential diagnostic or vaccine targets. Finally, we leverage publicly deposited proteomics data from human cells infected with SARS-CoV-2, as well as a second study with the closely related MERS-CoV to identify the two proteins of highest abundance in human infections. The resulting final list contains the 24 peptides most unique and diagnostic of SARS-CoV-2 infections. These peptides represent the best targets for the development of antibodies are clinical diagnostics. To demonstrate one application of this we model peptide fragmentation using a deep learning tool to rapidly generate targeted LCMS assays and data processing method for detecting CoVID-19 infected patient samples.

**Graphical Abstract:** 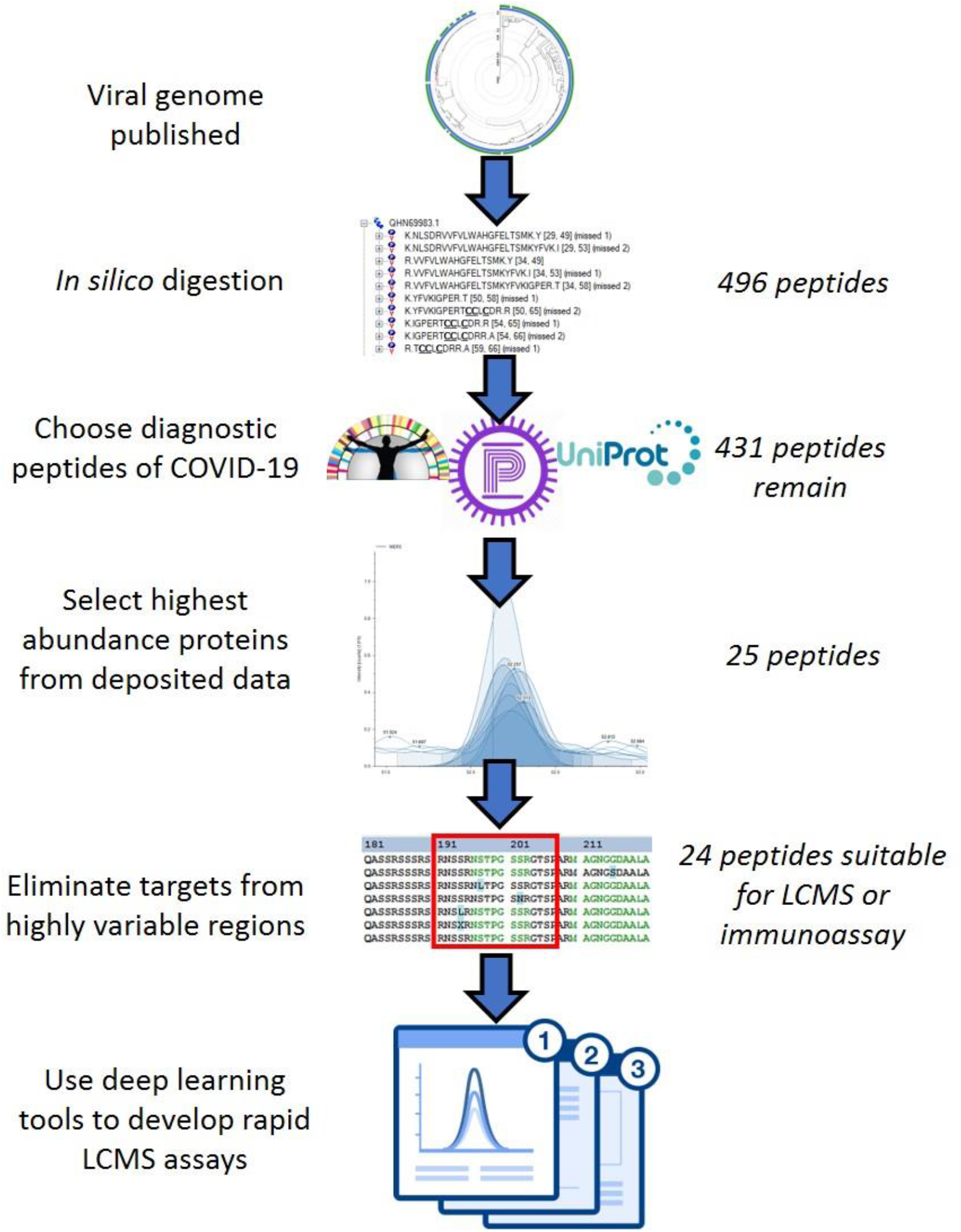

## Introduction

The identification of peptides expressed unique to pathogens is required for the development of diagnostic assays as well as vaccine targets. Antibody based techniques such as enzyme linked immunosorbent assay rely on antibodies raised against specific peptide targets. Mass spectrometry (MS) based diagnostic assays typically require many rounds of optimization to identify peptides that are unique in both sequence or in chemical characteristics to distinguish them from the complex host background.^1^

The detection of viral proteins in body fluids can be a rapid and specific diagnostic for infection in severe acute respiratory syndrome (SARS).^2–4^ During the 2003 (SARS) outbreak, non-MS based methods of protein detection proved to be more successful^5,6^ than LCMS methods.^7–9^ Non-MS based methods, such as western blots, enzyme-linked immunosorbent assays (ELISAs), and protein arrays, rely on antibodies for the detection of proteins. Given recent studies concerning high variability in antibody production, LCMS-based methods are an attractive alternative approach for the rapid identification of small molecules, proteins, and peptides in clinical settings where consistency is paramount.^10,11^

In the 15 years since the 2003 SARS outbreak, LCMS technology has experienced a revolution led primarily by increases in the speed, sensitivity, and resolution of MS instruments. Today, protein array and antibody-based methods are falling out of favor in both research and clinical diagnostics, due in large part to the improvements in LCMS technology.^12,13^ A review of this growth by Grebe and Singh described a clinical lab with no LCMS systems in 1998 that completed over 2 million individual LCMS clinical assays in 2010.^14^ Incremental improvements in rapid sample preparation techniques, chromatography, and data processing have also contributed to the increasing use of LCMS-based clinical testing. A 2013 study demonstrated the level of advance by identifying 4,000 yeast proteins in one hour of LCMS run time, identifying approximately 75 proteins/min at a rate 100 times faster than studies a decade prior.^15^

Targeted peptide-centric assays are advantageous when sensitivity is paramount over the quantity of identified targets. Targeted assays often rely on tandem MS with high speed, but relatively low accuracy, quadrupoles.^16^ Quadrupoles can be used to select ions for fragmentation with quantification of fragment ions by other quadrupoles in single reaction monitoring (SRM). They can also be used in conjunction with high resolution systems in single ion monitoring (SIM) and parallel reaction monitoring (PRM). SRM relies on fragmentation which requires *a priori* the mass to charge ratio (m/z) of both the peptide ion of interest and its dominant ions produced during fragmentation. Commonly, the collision energy is optimized for each peptide fragment in order to maximize efficiency and signal. SIM uses higher resolution scans in lieu of fragmentation, which requires *a priori* only an approximate m/z ratio for the peptide. In SIM targeted assays, the peptide ion’s exact mass is extracted post-run during data processing. Two studies have shown that SIM scans with ≥60,000 resolution produce selectivity comparable to SRM.^17,18^ PRM targeted assays combine fragmentation and high resolution molecule monitoring. In contrast to SIM, isolated ions are subjected to fragmentation. Post-run data processing then calculates quantification from the high-resolution accurate mass of the fragment ions. For proteins with only one available peptide target, PRM is the most selective technique in modern proteomics.^19,20^

Due to the high sensitivity of targeted LCMS assays, background signal is of central concern. Peptides must be selected that are unique in one or more of the following characteristics to be truly diagnostic; m/z, peptide sequence, or hydrophobicity. Due to the complexity of the human background thousands of peptides may be possible that are within 0.1 Da in m/z.^21^ In the case of ions with high m/z similarity, fragmentation to reveal peptides with sequences of low homology to the background can be used to easily discern these targets. The elution time of the peptides, once characterized and monitored with internal standards can be utilized to distinguish between even highly similar peptides, including sequences that are chemically modified but on alternative residues.^22^

To address these concerns an increasing number of in silico tools have been developed that can speed the identification of suitable LCMS targets. Tools such as Picky,^23^the Phosphopedia,^22^the Peptide Manager, and Purple^1^, and Skyline^24^ allow the removal of peptides from theoretical proteome backgrounds or the intelligent selection of peptide targets based on other parameters. Peptide retention times can be calculated based on amino acid hydrophobicity calculations, although the inclusion of internal reference calibration standards such as ProCal are necessary for adjustments to different chromatographic conditions.^25^

In traditional LCMS targeted assay development, once suitable targets are identified, peptides are then synthesized, and ideal fragment ions are selected for use following rounds of selection and optimization. One way to improve development time is by using databases of experimental fragmentation data such as the PeptideAtlas.^26^ However, peptide fragmentation follows specific energetic patterns, resulting primarily in fragments caused by separation at the peptide bond. It is therefore possible to create theoretical spectral libraries *in silico* from peptide sequence alone. Theoretical spectral libraries are especially useful when the biological samples are unavailable. New tools that employ deep learning algorithms have been demonstrated to produce theoretical MS2 spectra superior to previous prediction models and, in the absence of true experimental data, are the best resources currently available.^27,28^ These deep learning algorithms can learn from vast libraries of experimental data to predict the fragmentation patterns of new peptide sequences that they are given. One such algorithm, PROSIT, uses the vast synthetic human peptide libraries, from the ProteomeTools project^29^ for its training dataset. Due to the high quality of the 450,000 synthetic peptides experimentally fragmented in ProteomeTools to date, PROSIT has been demonstrated to create spectral libraries that are, in some cases, superior to experimentally derived in-house spectral libraries.^30^

In this study, we describe a *in silico* approach to identify unique peptides presented during viral infection of human cells. Using a stepwise elimination approach, we outline strategies to begin with a theoretical protein sequence and remove peptides with homology in clinically relevant systems. By removing peptides with homology to human and the relevant human microbiome, we can eliminate many targets that would produce false positive tests for the virus in any healthy person. We further improve on this by removing peptides that could be present in diseases with a similar clinical presentation. By utilizing the data in public repositories, we can determine the protein targets with the highest abundance as well as the highest likelihood of being diagnostic in different assays. We further apply this method toward the analysis of SARS-CoV-2 infected materials by identifying regions of variability in field isolates identified to date and flag these peptides, as diagnostics targeted on these peptides may lead to false negatives. The restricted target list may be utilized toward the generation of immunological based assays such as ELISA or immunoswabs^2^ or provide peptides of value as vaccine targets.

To demonstrate an application of this approach we use the peptides identified in this work to create materials to facilitate LCMS studies of SARS-CoV-2 infected tissues. We provide along with this work necessary resources for both DDA and DIA investigation of these materials through the production of spectral libraries and a list of predicted PTMs investigators should consider when analyzing these materials. The peptide spectral libraries are further utilized to create transition lists optimized for hardware from 5 LCMS instrument vendors. All materials and methods described herein are available as supplemental material to this publication. The supplemental material includes protein sequences (FASTA files), predicted PTMs, theoretical MS2 spectral libraries, instrument methods and targeted method data processing templates.

Our work demonstrates not only the feasibility of this approach, but also its ability to rapidly develop methods even in the face of limitation of access to sample experimental data. We use the example of SARS-CoV-2 viral protein detection to underscore the power of today’s protein informatics tools in responding to an urgent public health crisis.

## Results and Conclusions

### Comparison of LCMS to Alternative Pathogen Detection Methods

Genetics based assays that rely on the polymerase chain reaction (PCR) have well established sensitivity due to the ability to amplify nucleotide chains. Primer design must be optimized and may be adversely affected by unknown complex matrices such as in human stool where the true depth of the microbiome may be unknown.^39^ PCR-based assays have also been considered cost-prohibitive in some areas of clinical microbiology compared to other assays.^40^ A comparison of the relative sensitivity of PCR and protein based assays is challenging due to the differences in nomenclature and in the actual targets being analyzed. A comparison of detection limits of real-time (RT) PCR and other methods for *Giardia* pathogens in human stool determined revealed conflicting results. When compared to a rapid immunoassay for *Giardia lambia*, PCR was found inferior to both a rapid immunoassay and manual microscopic analysis. A later similar work compared assays for *Giardia intestinalis* and concluded that RT-PCR was a superior method to ELISA or microscopic analysis. In addressing the detection of emerging pathogens PCR requires several steps to be deployed *en masse*. First, a genomic sequence must be generated from which primers can be designed. These primers must be evaluated for cross-reactions in clinical samples and these reagents must be generated in bulk, subjected to quality control (QC) procedures and shipped to the site of testing. The lead time for these reagents has been well-publicized in addressing the COVID-19 pandemic, as has the challenges in quality of the primers for an assay deployed by the US CDC in February, 2020.^41^ A summary of relevant assays is shown in Table 1. ELISA based assays for SARS-CoV display the least sensitivity of the these assays with detections in nanograms/mL.^3^ ImmunoSwab assays first described for SARS-CoV and recently shown to provide rapid utility for SARS-CoV-2 report sensitivity in the former case of 10 picograms/mL.^2^ Due to the high similarity of the two platforms we hypothesized that the SARS-CoV-2 assay will be of similar performance. Two examples of optimized peptide pharmacokinetics assays in serum have demonstrated the utility of rapid LCMS based targeted assays that can achieve limits of detection and assay time comparable to current validated assays.

**Table 1.** A comparison of relevant assays and their respective limits of detection.

| Assay and Reference | Matrix | Detection limit | Assay Time |
| --- | --- | --- | --- |
| SARS-CoV Nucleocapsid ImmunoSwab<sup>2</sup> | Nasopharyngeal aspirate medium | 10 picogram/mL | 45 min |
| SARS-CoV-2 Nucleocapsid Immunoswab<sup>42</sup> | Nasopharyngeal aspirate/urine | Unstated / based on assay above | 10 min |
| SARS-CoV Nucleocapsid ELISA<sup>4</sup> | Nasopharyngeal aspirate | 2.5 nanogram/mL | 4-hour EST |
| SARS-CoV Nucleocapsid ELISA<sup>4</sup> | Stool | 9.0 nanograms/mL | 4-hour EST |
| SARS-CoV Protein Microarray<sup>43</sup> | Human Serum | 1:64,000 Dilution of Positive Serum | ~2 hours |
| Inflammatory Mediator Protein Microarray<sup>44</sup> | Human Serum | 20 picogram/mL | ~3 hours |
| CDC-SARS-CoV-2<sup>45</sup> | Nasopharyngeal aspirate | 3.2 copies/μL | 4 hours |
| SARS-CoV-2 RT-Lamp/CAS12<sup>45</sup> | Nasopharyngeal aspirate | 10 copies/μL | 45 min |
| Enkephalin LCMS<sup>46</sup> | Plasma | 10pg/mL | 6.5 min |
| Orexin LCMS<sup>47</sup> | CSF | 4pg/mL | 7 min |

### Theoretical peptides

In shotgun proteomics, proteins are first digested into smaller peptide fragments that are more easily detected by the instrument. Given its widespread use, high efficiency and speed of digestion, we chose to develop methods that exclusively use the proteolytic enzyme trypsin, which produces “tryptic peptides.” Sequencing grade trypsin exhibits high efficiency cleavage at unmodified (1) arginine and (2) lysine residues unless followed by a proline. Trypsin also has the advantage of leaving a terminal basic residue at the cleavage site, which increases the likelihood of complete fragment ion coverage from the charged terminal.^48^ Due to these reasons, trypsin is utilized unless the protein sequence has an abnormally high or low number of lysine or arginine residues. A very high frequency of the residues (such as Lysine-rich proteins) will create very short peptides that could be uninformative for protein identification. A very low frequency of the residues will create very large peptides, or undigested (“intact”) proteins in some cases, that are difficult to detect and fragment. Our theoretical trypsin digest of the 2019-nCoVpFASTA1 database produced tryptic peptides with average lengths of 8 to 18 amino acids. These results indicate that trypsin digestion is an appropriate choice for detection of these viral proteins.

### Removal of proteome theoretical background

The recently described program Purple was used for background subtraction of relevant theoretical proteomes using default parameters. Peptides were considered relevant with a sequence between 6 and 24 amino acids in length. Using these parameters, the COVID-19 UniProt FASTA produced 496 unique peptide sequences. In the first stage analysis we eliminated human peptides from further consideration. By subtracting all the human proteome background, we find that two peptide sequences are identical in SARS-CoV-2 and the human sequence. For example, the peptide sequence TLLSLR is found in the Replicase 1a protein of SARS-CoV-2 as well as in 4 human proteins, including Focadhesin (S4R400). These identical peptides should be excluded due from any further analysis due to these identical sequences. Purple was also used to remove peptides of identical amino acid composition with alternate sequence according the to default parameters previously described,^1^ removing another 45 peptides from this list.

In the second stage we developed a pipeline to remove proteins derived from the complex human microbiome from consideration. The NIH Microbiome initiative has created vast libraries using genomic sequencing technologies to identify the organisms present in and on *Homo sapiens*.^49,50^ These libraries do not, however, address protein abundance. In order to create a relevant proteome background for subtraction we performed a meta-analysis of a recent in depth study of the human saliva metaproteome.^38^ The raw instrument data was searched using the complete NIH saliva microbiome FASTA database of 536,284 protein sequences derived from 249 organisms as well as the complete UniProt catalog of reviewed bacterial and human protein sequences to date as well as the common list of protein lab contaminants. A new database was created of all protein sequences that contained at least one peptide spectral match (PSM), resulting in 29,816 unique protein sequences. Although it is possible that the proteins provided in the NIH FASTA are possible in the saliva microbiome, without adjusting for protein abundance and removing redundancy for shared proteins between organisms the proteome background may be misconstrued as more complex than it truly is. By removing from consideration, the 29,816 proteins of most relevant abundance we can more accurately measure the proteome background (data not shown.) Subtraction of the human microbiome background removed another 12 peptides that should be excluded in oral diagnostics or in the development of vaccine targets for SARS-CoV-2.

For the third stage subtraction we narrowed the peptide list to those targets that would be diagnostic for SARS-CoV-2 infection but would not cross-react with organisms of similar symptoms or clinical presentation. We created separate protein FASTA databases from UniProt 2020_01 for the viral pathogens, rhinovirus, influenza, pneumoniae, pneumophila and respiratory syncytial virus (RSV) as well as staph aureus and streptococcus species. The subtraction of these proteins from the background removed another six peptides from consideration.

The final stage of proteome background subtraction focused on the removal of peptides that could be derived from other coronavirus strains, including MERS, SARS-CoV and other protein sequences. Unsurprisingly, this proved to be the largest removal step, with 306 peptides subtracted from consideration and only 125 remaining as clearly unique and diagnostic of SARS-CoV-2. These results are available as Supplemental X and visualized in Figure 1.

**Figure 1.**
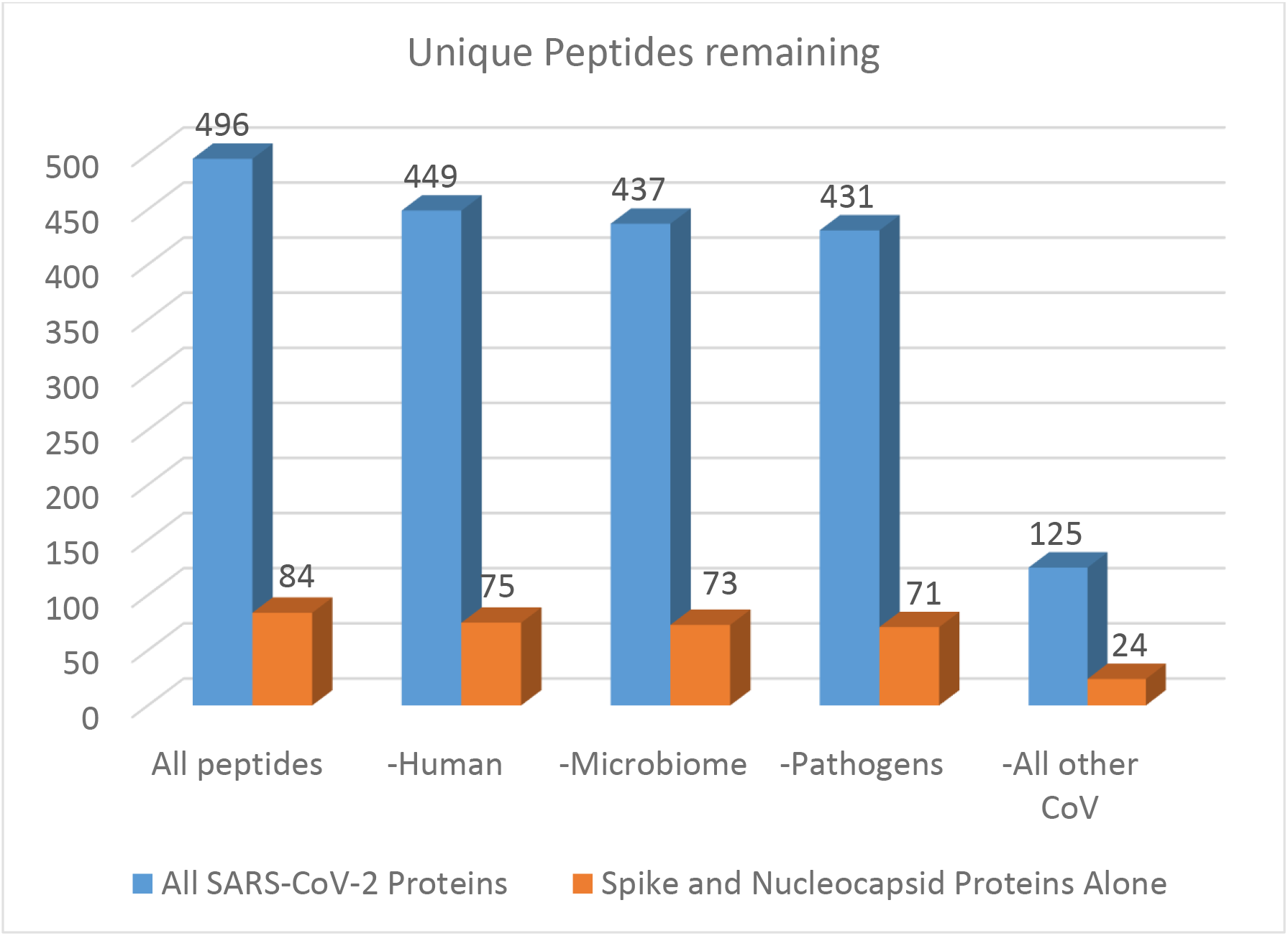
A summary of the peptide numbers as they are removed from an increasingly complex theoretical proteome background

### Utilization of Publicly deposited proteomics data to identify ideal protein targets

As shown in Table X, several studies for related coronavirus strains had been deposited in public repositories. In March of 2020, three new studies were released featuring work with SARS-CoV-2. In order to determine if some viral proteins would be of higher relative abundance in human infection, we first began by searching previously released, but unpublished, files from MERS-CoV infection of Calu3 cells. As shown in Figure 2A more peptides were identified from the nucleocapsid protein than any other protein, followed by the Spike protein. When the first SARS-CoV-2 study was released we reprocessed this study and observed the same trend, as shown in Figure 2B. Although not strictly quantitative, counting the number of peptides identified in a global proteomics experiment has been historically used as a metric for approximating relative protein abundance in a sample. These results suggest that the Nucleoprotein and Spike Glycoprotein may be the highest abundance SARS-CoV-2 proteins present in human infections

**Figure 2A.**
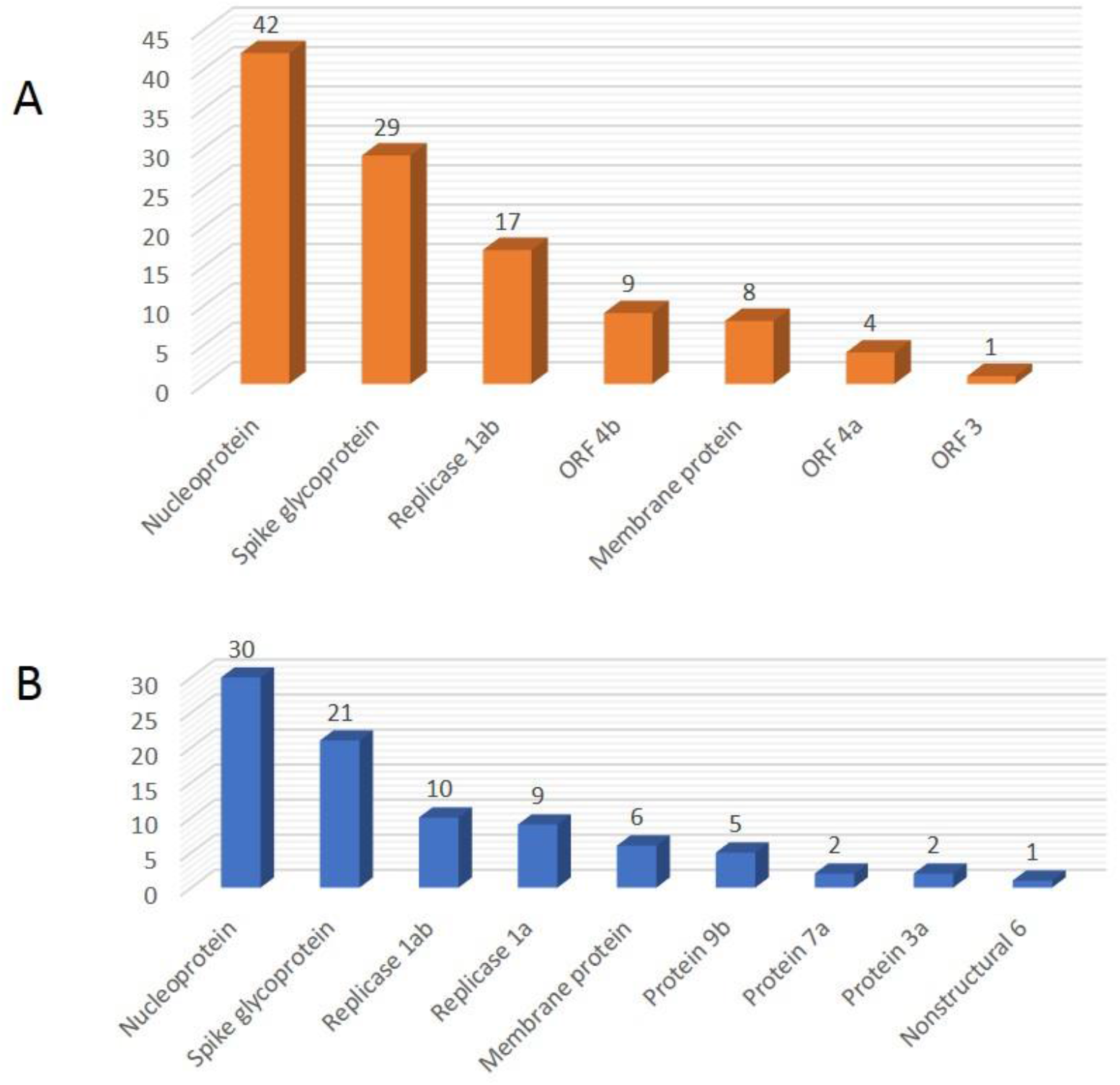
The number of unique peptides identified in a metanalysis of MERS-CoV infection for each protein in human cells. **2B**. The same analysis performed on the first proteomic study released on SARS-CoV infected human samples.

### Identification of peptides most likely to be affected by evolutionary pressures

To identify peptides likely to make poor targets due to genomic variability, we performed a 2-stage approach. The first consisted of aligning and visualizing protein sequences being deposited in NCBI MetAlign and AliView software in R. These tools allow the color coding and rapid flagging of regions of alternative protein sequences in those aligned. A visualization of one such region from the nucleocapsid protein is shown in Figure 3A. To verify these results, we processed proteomics data derived from cloned nucleocapsid proteins utilizing every unique full length nucleocapsid protein sequence available in UniProt. Figure 3B mirrors the results from the visualization analysis. We identify one peptide region in the nucleocapsid and two in the spike glycoprotein that may provide false positives for diagnostic assays.

**Figure 3A.**
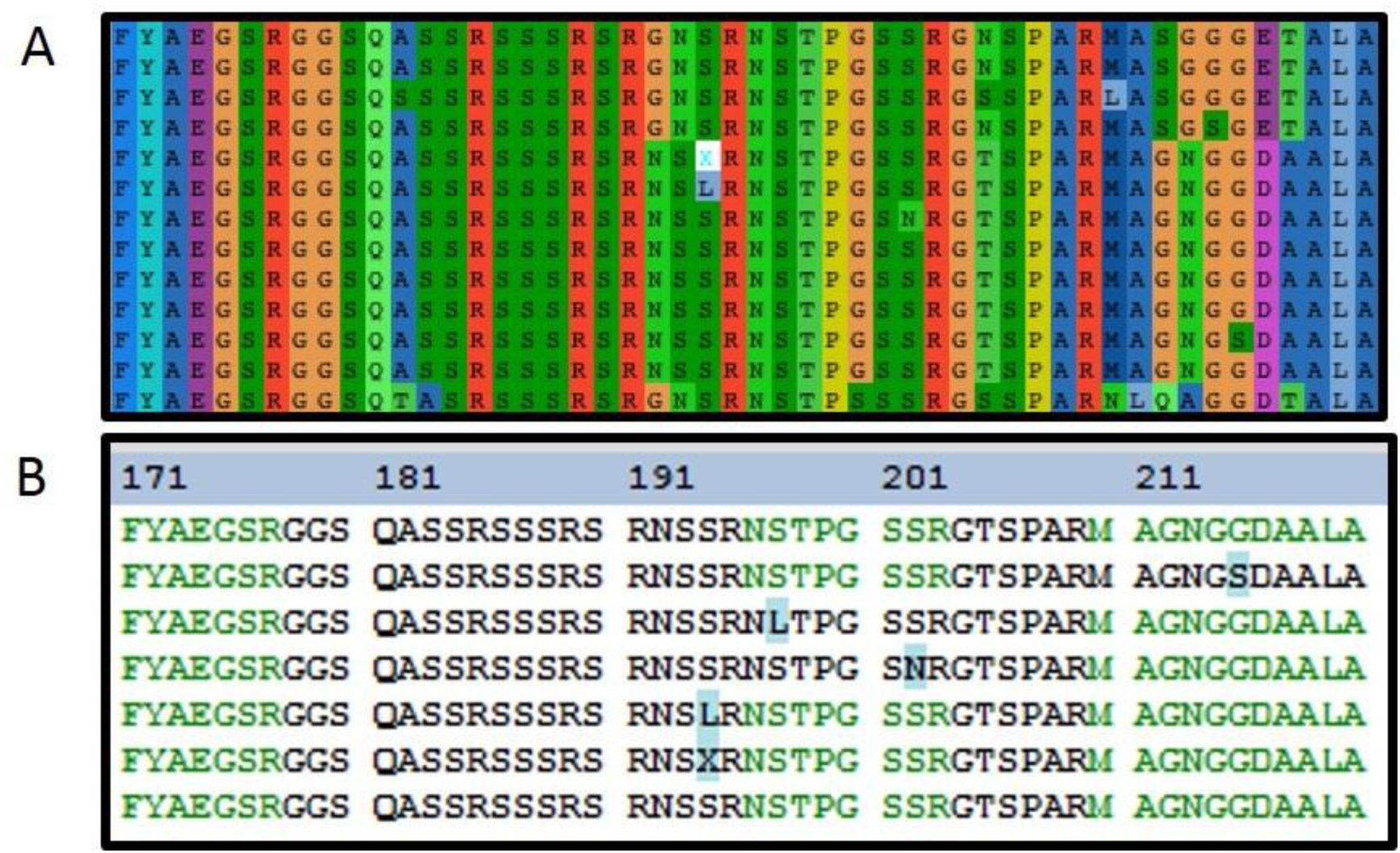
AliView output predicting protein variability in a region of the nucleocapid protein sequence. **3B**. Comparison of all available annotated NCBI protein sequences to recently deposited proteomics data that reflects the variability in this viral protein region.

### Generation of Spectral Libraries and Experimental Methods

In the absence of any experimentally derived spectra from SARS-CoV-2 proteins we used the Prosit deep learning tool to generate theoretical spectra for each peptide. Figure 4 is an example of a nucleocapsid peptide predicted by Prosit on January 27^th^ compared to the first experimental data available for a cloned protein from this virus. The top panel is the experimental spectra and the bottom is a mirror plot showing the predicted fragments and their expected relative intensities. Prosit accurately predicted the absence of fragments y14 and y15 as well as the presence of only b ions central to the peptide sequence. Furthermore, the abundance predictions allow for the selection of the most appropriate ions for MS/MS assays such as PRM and MRM. As one utility of the methods described herein, we used the Skyline software to create transition lists optimized specifically for each of the Skyline-compatible triple quadrupole instruments. While minor modifications are required for SCIEX, Agilent, and Shimadzu instruments, Waters Xevo and Thermo instruments use identical parameters for transition list design. Most modern triple quadrupole instruments are capable of 500 SRMS/sec and fully permit the use of 2,000 transition lists, as provided here. For older instruments that lack this scan speed or that require higher dwell times, the transition lists included in the supplemental methods may be reduced by the end user accordingly.

**Figure 4.**
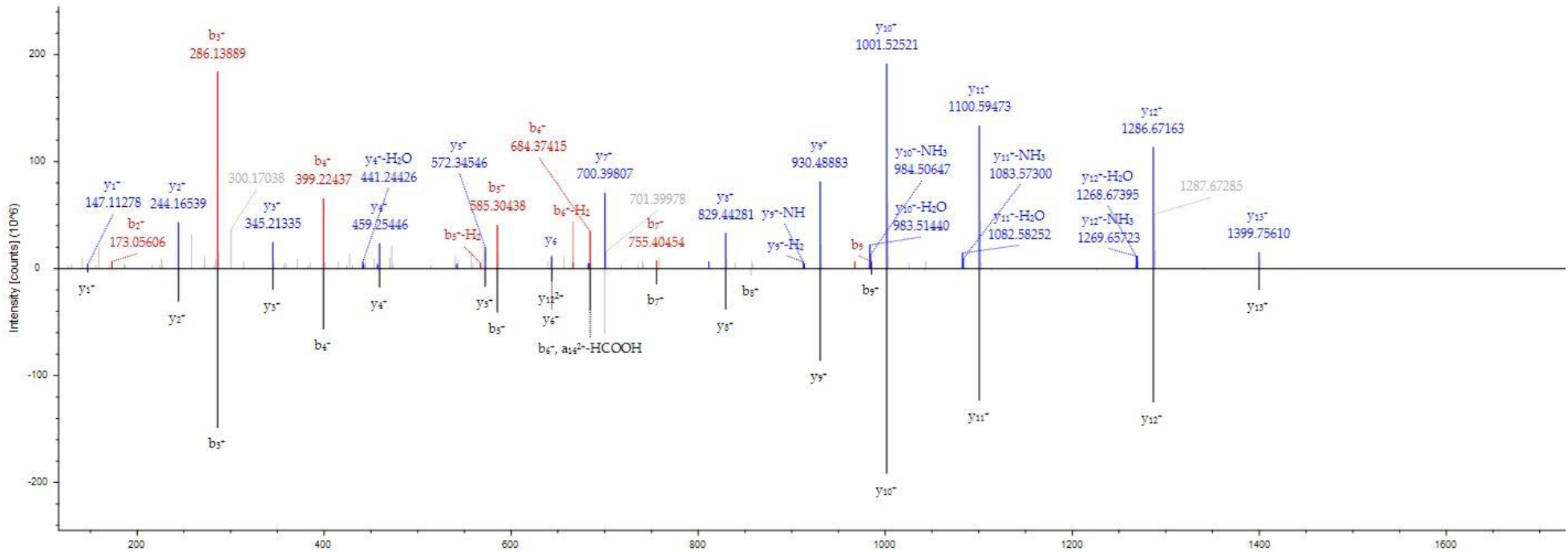
A mirror plot demonstrating the Prosit derived MS/MS spectra (bottom half) against the first known fragmentation pattern for this SARS-CoV-2 nucleocapsid peptide in the literature (top half)

PRM methods monitor multiple transitions simultaneously but at a time cost. The highest scan speed currently available in Orbitrap instruments is 48 scans per second and is only available on the Exploris 480 system (data not shown). In order to achieve maximum sensitivity, higher fill times are often required for these instruments. We chose to utilize three peptides/protein for these methods. Alternative peptides can be selected from the Skyline files provided (Supplemental Information) or by selecting peptide mass targets from the other transition lists. A summary of the resources provided in this work are available in the Supplemental Information.

### Resources for untargeted proteomics methods

In the course of this work, we created the minimal resources required for untargeted shotgun analysis of SARS-CoV-2 infected materials. These resources are made available in the supplemental material including the full Prosit Spectral libraries, FASTA files and predicted post-translational modifications. The Prosit spectral libraries (Supplemental Material) enable the interrogation of DIA data and may be used for DDA experiments that employ tools such as the MSPepSearch (NIST).^51^ DDA data requires only the protein FASTA file and a list of PTMs that may be present in the sample. Our analysis using ModPred predicted 17 possible PTMs (Supplemental Table 1). Amidation was the most frequent predicted PTM, but there is no known biological mechanism that we could derive from a survey of the literature. Palmitoylation, the second most frequent predicted PTM, is a well characterized viral PTM with critical functions in human immunodeficiency virus (HIV), human herpes virus (HHV), and influenza virus infectivity.^52–54^ During the assembly of this manuscript, proteomics data became available from the infection of *Chlorocebus sabaeus* (Green monkey) cells with SARS-CoV-2. Multiple phosphorylation sites observed on the nucleocapsid protein from this study were accurately predicted as high confidence sites by the ModPred software, including S176 and S205, further strengthening our confidence in this program for this function (Supplemental Table 1).

### LCMS Data for SARS-CoV-2

The COVID-19 global health crisis has permitted an unprecedented degree of scientific sharing and progress. Prior to March 11, 2020 no proteomics data existed in any public repository for SARS-CoV-2 infected materials. Table 2 is a summary of the LCMS data available as of the date of this writing for both SARS-CoV-2 and the most closely related coronavirus strains as recently described.^55^

**Table 2.**
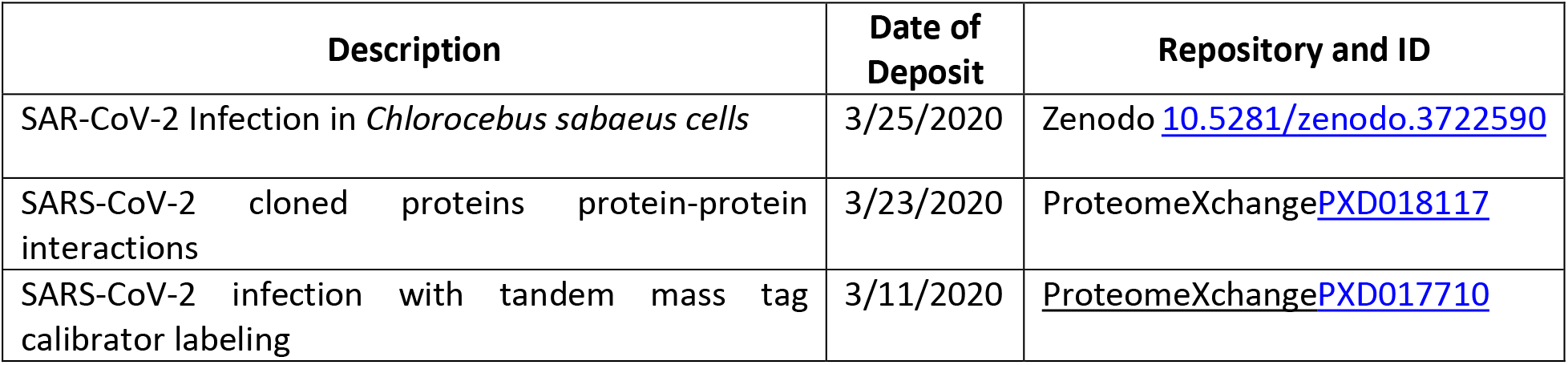
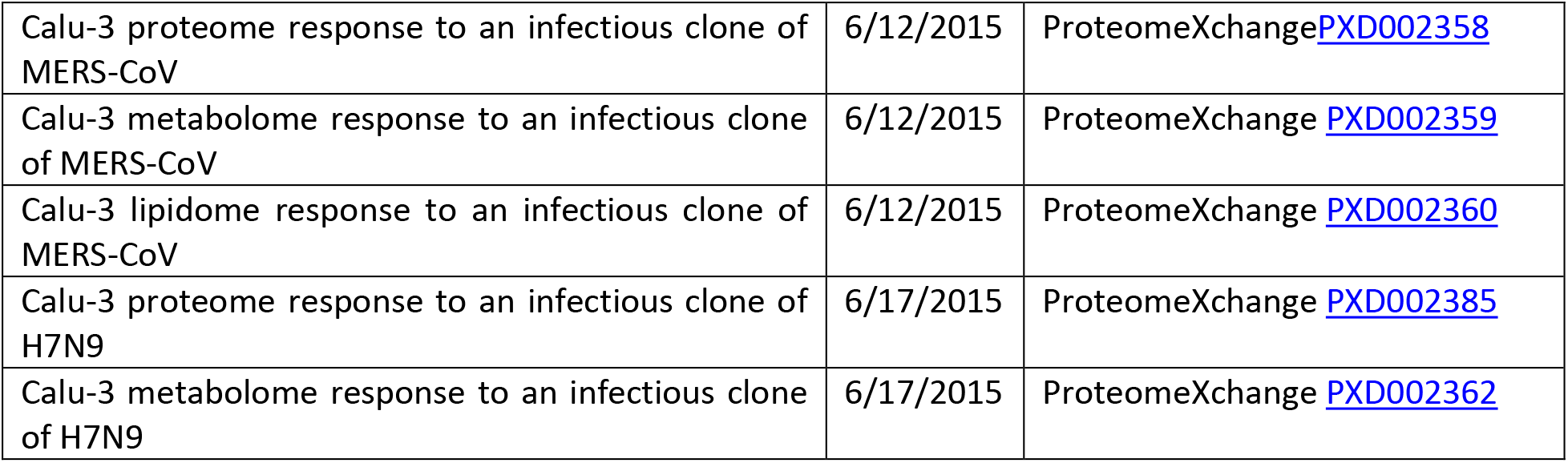
Publicly available data relevant to SARS-CoV-2 Studies

| Description | Date of Deposit | Repository and ID |
| --- | --- | --- |
| SAR-CoV-2 Infection in *Chlorocebus sabaeus* cells | 3/25/2020 | Zenodo [10.5281/zenodo.3722590](https://zenodo.org/record/3722590) |
| SARS-CoV-2 cloned proteins protein-protein interactions | 3/23/2020 | ProteomeXchange [PXD018117](https://proteomeexchange.org/data/PXD018117) |
| SARS-CoV-2 infection with tandem mass tag calibrator labeling | 3/11/2020 | ProteomeXchange [PXD017710](https://proteomeexchange.org/data/PXD017710) |
| Calu-3 proteome response to an infectious clone of MERS-CoV | 6/12/2015 | ProteomeXchange PXD002358 |
| Calu-3 metabolome response to an infectious clone of MERS-CoV | 6/12/2015 | ProteomeXchange PXD002359 |
| Calu-3 lipidome response to an infectious clone of MERS-CoV | 6/12/2015 | ProteomeXchange PXD002360 |
| Calu-3 proteome response to an infectious clone of H7N9 | 6/17/2015 | ProteomeXchange PXD002385 |
| Calu-3 metabolome response to an infectious clone of H7N9 | 6/17/2015 | ProteomeXchange PXD002362 |

## Summary

Using *in silico* methods, we have developed methods for the rapid selection of unique viral peptides for diagnostic assay and vaccine development, using the example of SARS-CoV-2. *In vitro* validation of these peptides and methods is required and outside the scope of this work given our current lack of access to such samples. To demonstrate the utility of our pipeline, we have provided the minimum materials for data processing for both DDA and DIA untargeted proteomics methods with FASTA databases, spectral libraries and by predicting relevant PTMs for consideration. To broaden the number of labs that can apply our methods, we optimized run parameters for widely used LCMS systems compatible with Skyline, representing instruments from five companies. We will continue to refine these resources and post updates to these methods to LCMSmethods.org and invite researchers anywhere in the world to contact us for assistance in further optimization to address this emerging threat.

## Acknowledgements

We thank Dr. Matthew Monroe at Pacific Northwest National Laboratory (PNNL) for assistance curating unpublished data in the ProteomeXchange/MASSIVE public repository.

## Author Contributions

Conceptualization, B.C.O and N.M.B.; Methodology, B.C.O. and C.J.; Investigation, S.M.M, C.J., B.A.N.; Writing-Original Draft, B.C.O, C.J. and N.M.B.; Writing-Review and Editing, B.C.O, B.A.M and N.M.B., Resources, C.J. and S.A.M.; Supervision, N.M.B. and B.C.O.

## Declaration of Interests

The authors declare no competing interests

## Methods Details

### Coronavirus FASTA databases

At the date of this writing, only theoretical protein sequences for SARS-CoV-2, are available. These sequences are being acquired and annotated and the result of translation of genomic sequence information. All sequences in this study were obtained from NCBI accession: txd2697049, https://www.ncbi.nlm.nih.gov/protein/?term=txid2697049). Using Proteome Discoverer 2.4 (Thermo), the protein sequences were combined into a single protein FASTA database (2019-nCOVpFASTA1; Supplemental Information), and added to human proteome sequences (UniProt SwissProt Human database; downloaded 2/15/2020) to produce a database including both human and COVID-19 protein sequences (Human_plus_2019-nCOVpFASTA2; Supplemental Information). The UniProt 2020_01 pre-released protein FASTA for SARS-CoV-2 was downloaded on 3/15/2020 is also provided here. (UniProt_202001_prerelease_COVID19. Fasta; Supplemental material)

### Publicly available proteomics data from human samples infected with other coronaviruses

Publicly available experiments on other coronavirus experiments were found by searching the ProteomeXchange Consortium web interface (http://www.proteomexchange.org/).^31,32^ Clarification of the identity of unpublished data from Pacific Northwest National Laboratory (PNNL) was provided by Dr. Michael Monroe. SARS-CoV-2 labeled proteomics data was made available ahead of publication^33^ as ProteomeXchange PXD017710. SARS-CoV-2 protein-protein interaction RAW files were downloaded from ProteomeXchange PXD018117.^34^

### Creation of peptide spectral libraries exclusive to SARS-CoV-2 with Prosit deep learning

The 2019-nCOVpFASTA1 (Supplemental Information) was converted to PROSIT peptide format with the EncyclopeDIA^35^ software, resulting in an *in silico* peptide digestion (parameters: Charge Range = 2-3; Max missed cleavage = 1, m/z range = 400-1,500; default NCE = 30eV; default charge = +2). PROSIT peptide fragmentation prediction libraries were generated using the PROSIT online web portal (https://www.proteomicsdb.org/prosit/) with the spectral library modeling interface (options: prediction model = Prosit_2019_intensity; iRT prediction model = Prosit_2019_irt). The libraries were exported in NIST MSP text format for use in Skyline.^24,36^

### PRM and SRM method development

For SRM transitions, the 2019-nCOVpFASTA1 (Supplemental Information) was imported into Skyline v20.1.0.28 (University of Washington) along with the PROSIT tryptic peptide spectral library. Peptide settings and transitions were optimized within Skyline to reflect the vendor optimization requirements. For Agilent systems, the 20ms default dwell time was selected for the transition settings. For SCIEX instruments, the same dwell time was utilized as well as automatic optimization of the declustering potential and compensation voltage from the transition settings menu. For Waters, Thermo, and Shimadzu systems, no further settings were required for transition list generation. All transition lists were exported as unscheduled 15min methods. For PRM methods, three peptides were selected for each protein due to the increased time per scan relative to SRM methods. Instrument-specific Skyline files are included in the Supplemental information as described in Table 1.

### Prediction of PTMs

PTMs were predicted for the 2019-nCOVpFASTA1 proteins (Supplemental Information) using the ModPred web interface (www.modpred.org; accessed 1/31/2020).^37^ All PTMs available at the date of analysis were selected as theoretical sites, and the Basic non evolutionary model was applied. ModPred ranks each PTM as high, medium, or low confidence as previously described.^37^ The ModPred web interface can accept a maximum of 5,000 amino acids. In order to analyze YP_009724389.1 the predicted sequence had to be divided into five sequences, using a 100 amino acid overlap to avoid disrupting potential large motifs. The results of ModPred were compiled into a single spreadsheet with all modifications of all confidence levels. A second sheet was created that contained only the high confidence PTMs predicted, as well as a final summary for the counting of predicted high confidence PTM occurrence, provided as Supplemental Material as detailed in Supplemental Table 1.

### Evaluation of genomic stability in SARS-COV-2

To ensure that our assay targets did not lie in a mutable portion of the genome, all available SARS-CoV-2 genome constructs (as of March 21st, 2020) were downloaded from NCBI and aligned using MAFFT (sorted fasta output and automatic input detection; ver 7.453) and visualized with AliView (ver 1.26). The protein sequences for the nucleocapsid, polymerase, spike protein, and membrane glycoproteins were collected from NCBI’s IPG database. Search terms included: nucleocapsid, N protein, E protein, envelope, polymerase, replicase, rdrp, spike protein, surface protein, S protein. Entries labeled ‘hypothetical’ were not included. Final alignments visualized included only entries originating directly from the virus (excluding things such as Bat-SARS-CoV, civet, Rhinolophus, etc.). All collected accession numbers can be found in Supplemental File 1. The evaluation of suitable targets was performed by manual review.

### Evaluation of Nucleocapsid Protein Stability and Identification of Variable Regions

LCMS RAW instrument files from Gordon *et al.,*^34^ were downloaded from ProteomeXchange (Table 2). The files for cloned nucleocapsid protein were processed in PD 2.2 using the default template for Q Exactive Basic ID. The files were searched against the cRAP database, the UniProt 2020_01 FASTA and a compiled database of all full length nucleocapsid protein variants available in NCBI on 3/24/2020. Of the 15 protein sequences available, only 10 were flagged as having unique sequences by PD. The search results were not merged into protein groups in order to facilitate the visualization of protein sequences using the sequence comparison view post processing tool.

### Creation of saliva microbiome background interference databases

Files deposited in ProteomeXchange with identifier PXD004319 were utilized to create a saliva microbiome background.^38^ The files were searched in Proteome Discoverer 2.2 using the parameters detailed in Supplemental Table 1. The FASTA databases used were the complete Oral Microbiome.FA file from the NIH Microbiome initiative, a custom database of all bacterial species in UniProt 2020_01 as well as the Homo sapiens FASTA from the same, as well as the cRAP database of common lab protein and peptide contaminants. The database marker node was utilized to flag sequences derived from each FASTA. Protein grouping was not employed in this analysis to allow the largest possible sequence variation. All protein sequences with more than one potential peptide spectral match (PSM) were retained and exported. The FASTA database utilities tool in PD 2.2 was used to compile these results into a FASTA that contained only proteins in UniProt standard accession format, entitled Adjusted_Human_Saliva_Microbiome_3_24_20.FASTA (Supplemental Table 1).

### FASTA databases of pathogens with similar clinical presentation

Protein FASTA databases were downloaded from UniProt 2020_01 to create databases of proteins indicative of pathogens with similar clinical presentation using the following search terms: Coronavirus reviewed [no], Influenza, Middle East Respiratory, Pneumoniae, Respiratory Syncytial Virus, Rhinovirus, Staphylococcus aureus, Streptococcus reviewed [yes].

### Automatic removal of interfering peptide targets with Purple

To remove peptides that with homology to those of the COVID-19 theoretical peptides, Purple_portable_0.4.2 for Windows was installed locally. The UniProt 2020_01 release of the SARS-CoV-2 FASTA was placed into the same local file with the FASTA databases from Human and the FASTA databases of pathogens with similar clinical presentation. Two analyses were performed, the first to remove peptides that occur in human alone for use in serum or plasma-based LCMS assays. A second analysis was performed using these in addition to the saliva microbiome peptides for use in saliva or nasopharyngeal sampling methods.

### Evaluation of SARS-CoV-2 Proteomics Data for Evaluating In silico targets

Labeled proteomics of SARS-CoV-2 infected human cells were recently deposited as ProteomeXchange PXD017710. To evaluate the relative abundance of proteins during infectivity, the RAW files were reprocessed in PD 2.2, using the default workflow for TMT 10-plex quantification with the FASTA databases for UniProt human and the SARS-CoV-2 FASTA prerelease from UniProt 2020_01. Protein and peptide assemblies were manually reviewed to cross-reference suitable peptide targets.

### Data and Code Availability

All data described in this manuscript, including protein databases, instrument methods, transition lists and Skyline templates are included in the Supplemental Information as detailed in Supplemental Table 1.

## List of Supplemental Materials

Supplemental Table 1: LCMS resources provided in Zip File with this Submission

Supplemental Table 2: Peptide lists remaining following subtraction of each proteome background

Supplemental File 1: Alignments of Nucleocapsid and Spike Protein Variants and analysis details

Supplemental Material 1: Zip file containing all resources detailed in Supplemental Table 1

**Supplemental Table 1.**
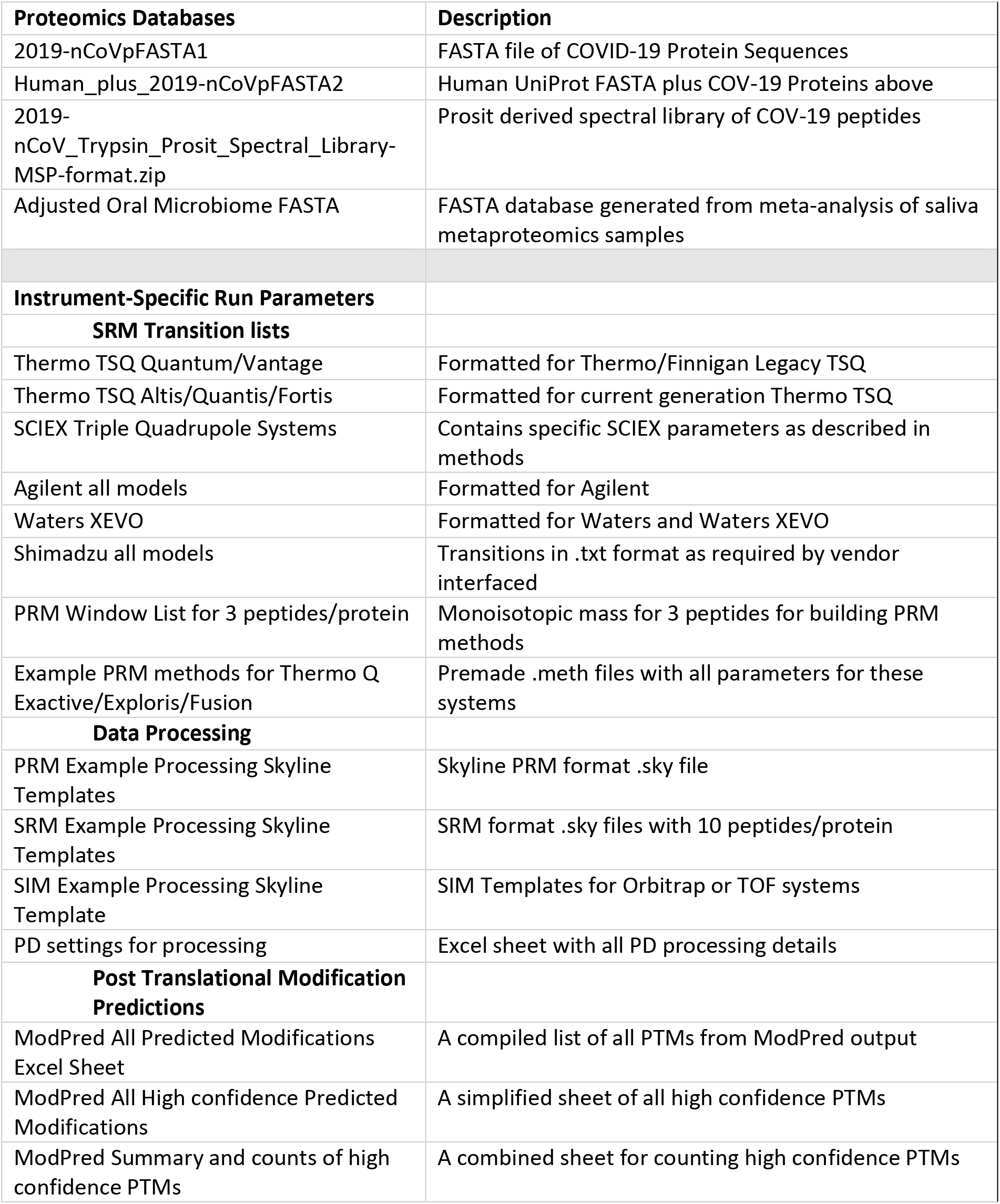
LCMS resources provided as supplemental material with this submission.

## Notes

https://www.lcmsmethods.org/

### Competing Interest Statement

The authors have declared no competing interest.

### Summary of Updates

Further work is presented here to select superior peptide targets for SARS-CoV-2 proteins. Peptides are excluded with identical or highly similar sequences to those from Human, the human saliva microbiome, as well as from diseases with similar clinical presentation. Furthermore, we highlight peptides from highly variable regions in the SARS-CoV-2 genomes that may lead to false negatives in diagnostic assays. Resources have been updated to reflect these changes

