## Supplemental Table 1 for "*In silico* approach toward the identification of unique peptides from viral protein infection: Application to COVID-19"

**Supplemental Table 1.** LCMS resources provided as supplemental material with this submission.

| **Proteomics Databases** | **Description** |
| --- | --- |
| 2019-nCoVpFASTA1 | FASTA file of COVID-19 Protein Sequences |
| Human_plus_2019-nCoVpFASTA2 | Human UniProt FASTA plus COV-19 Proteins above |
| 2019-nCoV_Trypsin_Prosit_Spectral_Library-MSP-format.zip | Prosit derived spectral library of COV-19 peptides |
| Adjusted Oral Microbiome FASTA | FASTA database generated from meta-analysis of saliva metaproteomics samples |
| **Instrument-Specific Run Parameters** |  |
| **SRM Transition lists** |  |
| Thermo TSQ Quantum/Vantage | Formatted for Thermo/Finnigan Legacy TSQ |
| Thermo TSQ Altis/Quantis/Fortis | Formatted for current generation Thermo TSQ |
| SCIEX Triple Quadrupole Systems | Contains specific SCIEX parameters as described in methods |
| Agilent all models | Formatted for Agilent |
| Waters XEVO | Formatted for Waters and Waters XEVO |
| Shimadzu all models | Transitions in .txt format as required by vendor interfaced |
| PRM Window List for 3 peptides/protein | Monoisotopic mass for 3 peptides for building PRM methods |
| Example PRM methods for Thermo Q Exactive/Exploris/Fusion | Premade .meth files with all parameters for these systems |
| **Data Processing** |  |
| PRM Example Processing Skyline Templates | Skyline PRM format .sky file |
| SRM Example Processing Skyline Templates | SRM format .sky files with 10 peptides/protein |
| SIM Example Processing Skyline Template | SIM Templates for Orbitrap or TOF systems |
| PD settings for processing | Excel sheet with all PD processing details |
| **Post Translational Modification Predictions** |  |
| ModPred All Predicted Modifications Excel Sheet | A compiled list of all PTMs from ModPred output |
| ModPred All High confidence Predicted Modifications | A simplified sheet of all high confidence PTMs |
| ModPred Summary and counts of high confidence PTMs | A combined sheet for counting high confidence PTMs |
