## Supplemental File 1 for "*In silico* approach toward the identification of unique peptides from viral protein infection: Application to COVID-19"

***Supplemental File 1. Alignment of Protein Variants***

***Part 1:*** Genome construct and protein accession numbers.

|  | **Accession Number** | **Description** |
| --- | --- | --- |
| **Genome Constructs** | MT072688 | Severe acute respiratory syndrome coronavirus 2 isolate SARS0CoV-2/61-TW/human/2020/ NPL, complete genome |
|  | MN988668 | Severe acute respiratory syndrome coronavirus 2 isolate 2019-nCoV WHU01, complete genome |
|  | NC_045512 | Severe acute respiratory syndrome coronavirus 2 isolate Wuhan-Hu-1, complete genome |
|  | MN938384 | Severe acute respiratory syndrome coronavirus 2 isolate 2019-nCoV_HKU-SZ-002a_2020, complete genome |
|  | MN975262 | Severe acute respiratory syndrome coronavirus 2 isolate 2019-nCoV_HKU-SZ-005b_2020, complete genome |
|  | MN985325 | Severe acute respiratory syndrome coronavirus 2 isolate 2019-nCoV/USA-WA1/2020, complete genome |
|  | MN988713 | Severe acute respiratory syndrome coronavirus 2 isolate 2019-nCoV/USA-IL1/2020, complete genome |
|  | MN994467 | Severe acute respiratory syndrome coronavirus 2 isolate 2019-nCoV/USA-CA1/2020, complete genome |
|  | MN994468 | Severe acute respiratory syndrome coronavirus 2 isolate 2019-nCoV/USA-CA2/2020, complete genome |
|  | MN997409 | Severe acute respiratory syndrome coronavirus 2 isolate 2019-nCoV/USA-AZ1/2020, complete genome |
|  | MN908947 | Severe acute respiratory syndrome coronavirus 2 isolate Wuhan-Hu-1, complete genome |
|  | MN988669 | Severe acute respiratory syndrome coronavirus 2 isolate 2019-nCoV WHU02, complete genome |
| **Nucleocapsid** | AAP30714.1 | putative nucleocapsid protein [SARS coronavirus CUHK-Su10] |
|  | AAR87518.1 | putative nucleocapsid protein N [SARS coronavirus TW11] |
|  | AAR12990.1 | nucleocapsid protein [SARS coronavirus HB] |
|  | AAS48456.1 | nucleocapsid protein [SARS coronavirus BJ01] |
|  | AAT76155.1 | nucleocapsid protein [SARS coronavirus TJF] |
|  | AAV49739.1 | nucleocapsid protein [SARS coronavirus B039] |
|  | AAU04673.1 | nucleocapsid protein [SARS coronavirus civet020] |
|  | AAU04658.1 | nucleocapsid protein [SARS coronavirus civet010] |
|  | AAV91640.1 | nucleocapsid protein [SARS coronavirus A022] |
|  | ACB69912.1 | nucleocapsid protein [SARS coronavirus BJ182-12] |
|  | AAX16200.1 | nucleocapsid protein N [SARS coronavirus WH20] |
|  | AAP50495.1 | nucleocapsid protein [SARS coronavirus FRA] |
|  | ACU31039.1 | nucleocapsid protein [SARS coronavirus Rs_672/2006] |
|  | ACZ71986.1 | nucleocapsid protein [SARS coronavirus ExoN1] |
|  | ACZ72117.1 | nucleocapsid protein [SARS coronavirus ExoN1] |
|  | ACZ72205.1 | nucleocapsid protein [SARS coronavirus ExoN1] |
|  | AAP82974.1 | nucleocapsid protein [SARS coronavirus Shanhgai LY] |
|  | AAS48575.1 | nucleocapsid protein, partial [SARS coronavirus xw002] |
|  | AAS48577.1 | nucleocapsid protein, partial [SARS coronavirus cw037] |
|  | AAS48576.1 | nucleocapsid protein, partial [SARS coronavirus cw049] |
|  | ARO76389.1 | nucleocapsid protein [Severe acute respiratory syndrome-related coronavirus] |
|  | NP_828858.1 | nucleocapsid protein [Severe acute respiratory syndrome-related coronavirus] |
|  | Q3LZX4.1 | Nucleocapsid protein |
|  | Q3I5I7.1 | Nucleocapsid protein |
|  | Q0Q468.1 | Nucleocapsid protein |
|  | QDF43838.1 | Nucleocapsid protein |
|  | NP_828858.1 | nucleocapsid protein [Severe acute respiratory syndrome-related coronavirus] |
|  | QHO62884.1 | nucleocapsid phosphoprotein [Severe acute respiratory syndrome coronavirus 2] |
|  | QHW06046.1 | nucleocapsid phosphoprotein [Severe acute respiratory syndrome coronavirus 2] |
|  | BCA37476.1 | nucleocapsid phosphoprotein, partial [Severe acute respiratory syndrome coronavirus 2] |
|  | QIC50514.1 | nucleocapsid phosphoprotein [Severe acute respiratory syndrome coronavirus 2] |
|  | BCB15098.1 | nucleocapsid phosphoprotein [Severe acute respiratory syndrome coronavirus 2] |
|  | QII57305.1 | nucleocapsid phosphoprotein [Severe acute respiratory syndrome coronavirus 2] |
|  | QII87775.1 | nucleocapsid protein, partial [Severe acute respiratory syndrome coronavirus 2] |
|  | QII87776.1 | nucleocapsid protein, partial [Severe acute respiratory syndrome coronavirus 2] |
|  | QIJ96530.1 | nucleocapsid phosphoprotein [Severe acute respiratory syndrome coronavirus 2] |
|  | YP_009724397.2 | YP_009724397.2 nucleocapsid phosphoprotein [Severe acute respiratory syndrome coronavirus 2] |
|  | QIK02783.1 | nucleocapsid phosphoprotein, partial [Severe acute respiratory syndrome coronavirus 2] |
|  | QIK02784.1 | nucleocapsid phosphoprotein, partial [Severe acute respiratory syndrome coronavirus 2] |
|  | QIH45050.1 | N protein [Severe acute respiratory syndrome coronavirus 2] |
|  | APO40586.1 | N protein [Severe acute respiratory syndrome-related coronavirus] |
|  | YP_009724392.1 | envelope protein [Severe acute respiratory syndrome coronavirus 2] |
|  | NP_828854.1 | protein E [Severe acute respiratory syndrome-related coronavirus] |
|  | APO40581.1 | E protein [Severe acute respiratory syndrome-related coronavirus] |
|  | AAS44718.1 | small envelope E protein [SARS coronavirus TW-GD1] |
|  | AAP51230.1 | envelope protein E [SARS coronavirus GD01] |
|  | ACB69908.1 | envelope protein E [SARS coronavirus BJ182-12] |
| **Polymerase** | YP_009725307.1 | RNA-dependent RNA polymerase [Severe acute respiratory syndrome coronavirus 2] |
|  | P0C6T7.1 | RNA-dependent RNA polymerase [Severe acute respiratory syndrome coronavirus 2] |
|  | ATU80197.1 | RNA-dependent RNA polymerase, partial [SARS-related betacoronavirus Rp3/2004] |
|  | ATU80158.1 | RNA-dependent RNA polymerase, partial [SARS-related betacoronavirus Rp3/2004] |
|  | ATU80102.1 | RNA-dependent RNA polymerase, partial [Bat SARS coronavirus HKU3] |
|  | ATU80112.1 | RNA-dependent RNA polymerase, partial [SARS-related betacoronavirus Rp3/2004] |
|  | ASB51350.1 | RNA-dependent RNA polymerase, partial [Severe acute respiratory syndrome-related coronavirus] |
|  | ASB51351.1 | RNA-dependent RNA polymerase, partial [Severe acute respiratory syndrome-related coronavirus] |
|  | AAQ18568.1 | RNA polymerase, partial [SARS coronavirus BJ2232] |
|  | AAP31512.1 | RNA polymerase 1b, partial [SARS coronavirus Tor2] |
|  | AAP06763.1 | RNA-directed RNA polymerase, partial [SARS coronavirus Hong Kong/03/2003] |
|  | AAP04003.1 | polymerase, partial [SARS coronavirus Vietnam] |
|  | AAP04587.1 | RNA-directed RNA polymerase, partial [SARS coronavirus Taiwan] |
|  | ARO76394.1 | RdRP, partial [Severe acute respiratory syndrome-related coronavirus] |
|  | NP_828869.1 | nsp12-pp1ab (RdRp) [Severe acute respiratory syndrome-related coronavirus] |
|  | ARO76378.1 | RdRP, partial [Severe acute respiratory syndrome-related coronavirus] |
|  | ARO76377.1 | RdRP, partial [Severe acute respiratory syndrome-related coronavirus] |
|  | ARO76379.1 | RdRP, partial [Severe acute respiratory syndrome-related coronavirus] |
|  | ARO76393.1 | RdRP, partial [Severe acute respiratory syndrome-related coronavirus] |
|  | AAS48581.1 | replicase 1b, partial [SARS coronavirus sf098] |
|  | P0C6F8.1 | Replicase polyprotein 1a |
|  | P0C6F5.1 | Replicase polyprotein 1a |
|  | P0C6V9.1 | Replicase polyprotein 1a |
|  | P0C6W6.1 | Replicase polyprotein 1a |
|  | P0C6T7.1 | Replicase polyprotein 1a |
|  | P0C6W2.1 | Replicase polyprotein 1a |
|  | AAP41036.1 | replicase 1AB [Severe acute respiratory syndrome-related coronavirus] |
|  | AAS48573.1 | replicase 1b, partial [SARS coronavirus cw037] |
|  | AAS44774.1 | replicase 1B, partial [SARS coronavirus TW-GD1] |
|  | AAS44803.1 | replicase 1B, partial [SARS coronavirus TW-GD1] |
|  | AAS48574.1 | replicase 1b, partial [SARS coronavirus sf098] |
|  | AAS44839.1 | replicase 1A, partial [SARS coronavirus TW-GD1] |
|  | AAS44821.1 | replicase 1B, partial [SARS coronavirus TW-GD1] |
|  | AAS48172.1 | replicase 1AB, partial [SARS coronavirus sf099] |
|  | AAS48174.1 | replicase 1AB, partial [SARS coronavirus sf098] |
|  | AAS48173.1 | replicase 1AB, partial [SARS coronavirus cw049] |
|  | AAS44806.1 | replicase 1B, partial [SARS coronavirus TW-GD4] |
|  | AGT21077.1 | replicase polyprotein 1ab [SARS coronavirus ExoN1] |
|  | AGT21076.1 | replicase polyprotein 1a [SARS coronavirus ExoN1] |
|  | ACV88184.1 | replicase 1AB polyprotein [Severe acute respiratory syndrome-related coronavirus] |
|  | AAV91630.1 | replicase 1AB [SARS coronavirus A022] |
|  | AAV49729.1 | replicase 1AB [SARS coronavirus B039] |
|  | AAT76146.1 | replicase 1ab [SARS coronavirus TJF] |
| Spike Protein | P59594.1 | Spike glycoprotein |
|  | ACZ72035.1 | spike glycoprotein precursor [SARS coronavirus MA15] |
|  | AAP50485.1 | spike glycoprotein [SARS coronavirus FRA] |
|  | AAP30030.1 | spike glycoprotein S [SARS coronavirus BJ01] |
|  | AAP51227.1 | spike glycoprotein S [SARS coronavirus GD01] |
|  | AAX16192.1 | spike glycoprotein S [SARS coronavirus WH20] |
|  | ACB69905.1 | spike glycoprotein S [SARS coronavirus BJ182-12] |
|  | ACB69894.1 | spike glycoprotein S [SARS coronavirus BJ182-8] |
|  | ACB69883.1 | spike glycoprotein S [SARS coronavirus BJ182-4] |
|  | ACB69860.1 | spike glycoprotein S [SARS coronavirus BJ182a] |
|  | AAU81608.1 | S protein [SARS Coronavirus CDC#200301157] |
|  | AAR91586.1 | spike glycoprotein S [SARS coronavirus NS-1] |
| **Membrane Glycoprotein** | AAS48455.1 | membrane protein [SARS coronavirus BJ01] |
|  | YP_009724393.1 | membrane glycoprotein [Severe acute respiratory syndrome coronavirus 2] |
|  | QII87843.1 | membrane glycoprotein [Severe acute respiratory syndrome coronavirus 2] |
|  | QIG55988.1 | membrane glycoprotein [Severe acute respiratory syndrome-related coronavirus] |
|  | QHR93621.1 | membrane glycoprotein, partial [Severe acute respiratory syndrome coronavirus 2] |
|  | AVP78034.1 | membrane protein [Bat SARS-like coronavirus] |
|  | Q0Q472.1 | M protein; Matrix glycoprotein |
|  | Q3I5J2.1 | M protein; Matrix glycoprotein |
|  | Q3LZX9.1 | M protein; Matrix glycoprotein |
|  | NP_828855.1 | matrix protein [Severe acute respiratory syndrome-related coronavirus] |
|  | AYV99793.1 | membrane protein [SARS coronavirus Urbani] |
|  | AYV99779.1 | membrane protein [SARS coronavirus Urbani] |
|  | ACZ72039.1 | membrane protein [SARS coronavirus MA15] |
|  | ACZ71756.1 | membrane protein [SARS coronavirus ExoN1] |
|  | AIA62333.1 | membrane glycoprotein [BtRs-BetaCoV/YN2013] |
|  | AIA62323.1 | membrane glycoprotein [BtRs-BetaCoV/GX2013] |
|  | AAS44719.1 | membrane glycoprotein M, partial [SARS coronavirus TW-GD1] |
|  | AAS44739.1 | membrane glycoprotein M, partial [SARS coronavirus TW-YM1] |
|  | ACB69909.1 | membrane protein M [SARS coronavirus BJ182-12] |
|  | AFR58704.1 | membrane protein [Severe acute respiratory syndrome-related coronavirus] |
|  | AFR58690.1 | membrane protein [Severe acute respiratory syndrome-related coronavirus] |
|  | AEA11017.1 | membrane protein [SARS coronavirus ExoN1] |
|  | AAP50489.1 | membrane protein [SARS coronavirus FRA] |
|  | ADE34769.1 | membrane protein [SARS coronavirus Frankfurt 1] |
|  | BAE93405.1 | membrane protein [SARS coronavirus Frankfurt 1] |
|  | ACZ71980.1 | membrane protein [SARS coronavirus ExoN1] |

***Part 2. Alignment of Nucleocapsid Proteins***

Nucleocapsid Alignment View

Mar 22, 2020

Sequences included in this alignment

| AAP30714.1 | putative nucleocapsid protein [SARS coronavirus CUHK-Su10] |
| --- | --- |
| AAR87518.1 | putative nucleocapsid protein N [SARS coronavirus TW11] |
| AAR12990.1 | nucleocapsid protein [SARS coronavirus HB] |
| AAS48456.1 | nucleocapsid protein [SARS coronavirus BJ01] |
| AAT76155.1 | nucleocapsid protein [SARS coronavirus TJF] |
| AAV49739.1 | nucleocapsid protein [SARS coronavirus B039] |
| AAU04673.1 | nucleocapsid protein [SARS coronavirus civet020] |
| AAU04658.1 | nucleocapsid protein [SARS coronavirus civet010] |
| AAV91640.1 | nucleocapsid protein [SARS coronavirus A022] |
| ACB69912.1 | nucleocapsid protein [SARS coronavirus BJ182-12] |
| AAX16200.1 | nucleocapsid protein N [SARS coronavirus WH20] |
| AAP50495.1 | nucleocapsid protein [SARS coronavirus FRA] |
| ACU31039.1 | nucleocapsid protein [SARS coronavirus Rs_672/2006] |
| ACZ71986.1 | nucleocapsid protein [SARS coronavirus ExoN1] |
| ACZ72117.1 | nucleocapsid protein [SARS coronavirus ExoN1] |
| ACZ72205.1 | nucleocapsid protein [SARS coronavirus ExoN1] |
| AAP82974.1 | nucleocapsid protein [SARS coronavirus Shanhgai LY] |
| AAS48575.1 | nucleocapsid protein, partial [SARS coronavirus xw002] |
| AAS48577.1 | nucleocapsid protein, partial [SARS coronavirus cw037] |
| AAS48576.1 | nucleocapsid protein, partial [SARS coronavirus cw049] |
| ARO76389.1 | nucleocapsid protein [Severe acute respiratory syndrome-related coronavirus] |
| NP_828858.1 | nucleocapsid protein [Severe acute respiratory syndrome-related coronavirus] |
| Q3LZX4.1 | Nucleocapsid protein |
| Q3I5I7.1 | Nucleocapsid protein |
| Q0Q468.1 | Nucleocapsid protein |
| QDF43838.1 | Nucleocapsid protein |
| NP_828858.1 | nucleocapsid protein [Severe acute respiratory syndrome-related coronavirus] |
| QHO62884.1 | nucleocapsid phosphoprotein [Severe acute respiratory syndrome coronavirus 2] |
| QHW06046.1 | nucleocapsid phosphoprotein [Severe acute respiratory syndrome coronavirus 2] |
| BCA37476.1 | nucleocapsid phosphoprotein, partial [Severe acute respiratory syndrome coronavirus 2] |
| QIC50514.1 | nucleocapsid phosphoprotein [Severe acute respiratory syndrome coronavirus 2] |
| BCB15098.1 | nucleocapsid phosphoprotein [Severe acute respiratory syndrome coronavirus 2] |
| QII57305.1 | nucleocapsid phosphoprotein [Severe acute respiratory syndrome coronavirus 2] |
| QII87775.1 | nucleocapsid protein, partial [Severe acute respiratory syndrome coronavirus 2] |
| QII87776.1 | nucleocapsid protein, partial [Severe acute respiratory syndrome coronavirus 2] |
| QIJ96530.1 | nucleocapsid phosphoprotein [Severe acute respiratory syndrome coronavirus 2] |
| YP_009724397.2 | YP_009724397.2 nucleocapsid phosphoprotein [Severe acute respiratory syndrome coronavirus 2] |
| QIK02783.1 | nucleocapsid phosphoprotein, partial [Severe acute respiratory syndrome coronavirus 2] |
| QIK02784.1 | nucleocapsid phosphoprotein, partial [Severe acute respiratory syndrome coronavirus 2] |
| QIH45050.1 | N protein [Severe acute respiratory syndrome coronavirus 2] |
| APO40586.1 | N protein [Severe acute respiratory syndrome-related coronavirus] |
| YP_009724392.1 | envelope protein [Severe acute respiratory syndrome coronavirus 2] |
| NP_828854.1 | protein E [Severe acute respiratory syndrome-related coronavirus] |
| APO40581.1 | E protein [Severe acute respiratory syndrome-related coronavirus] |
| AAS44718.1 | small envelope E protein [SARS coronavirus TW-GD1] |
| AAP51230.1 | envelope protein E [SARS coronavirus GD01] |
| ACB69908.1 | envelope protein E [SARS coronavirus BJ182-12] |

duplicate sequence	partial sequences in ovals**

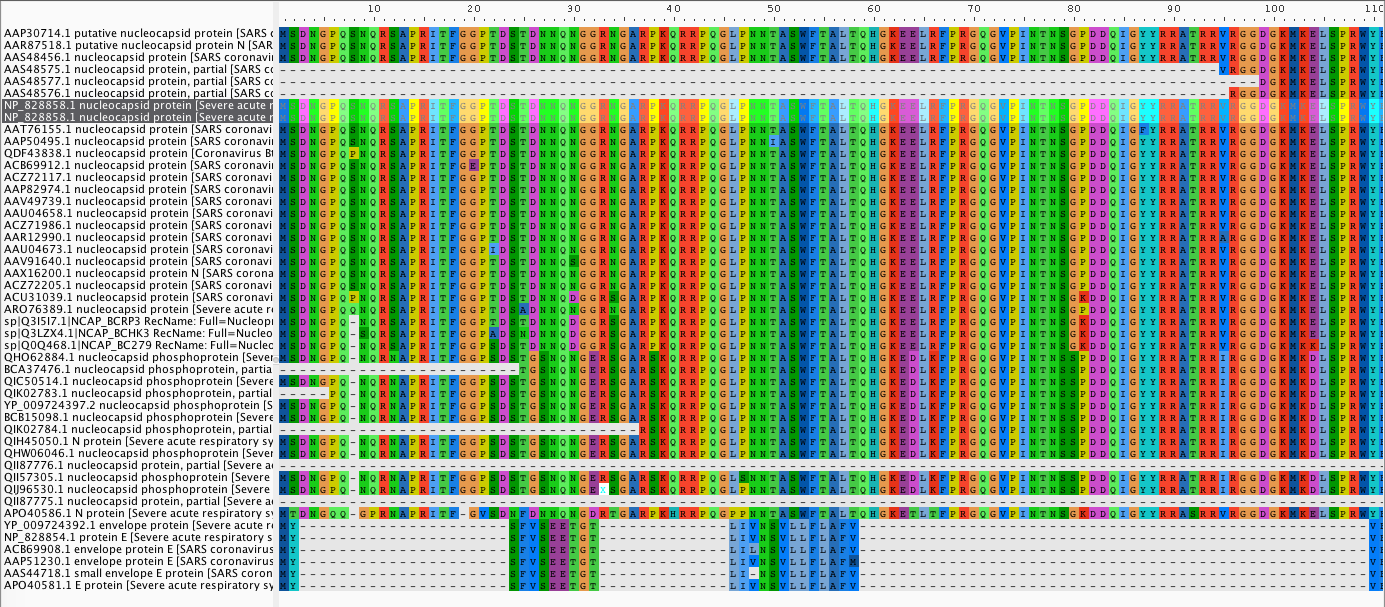

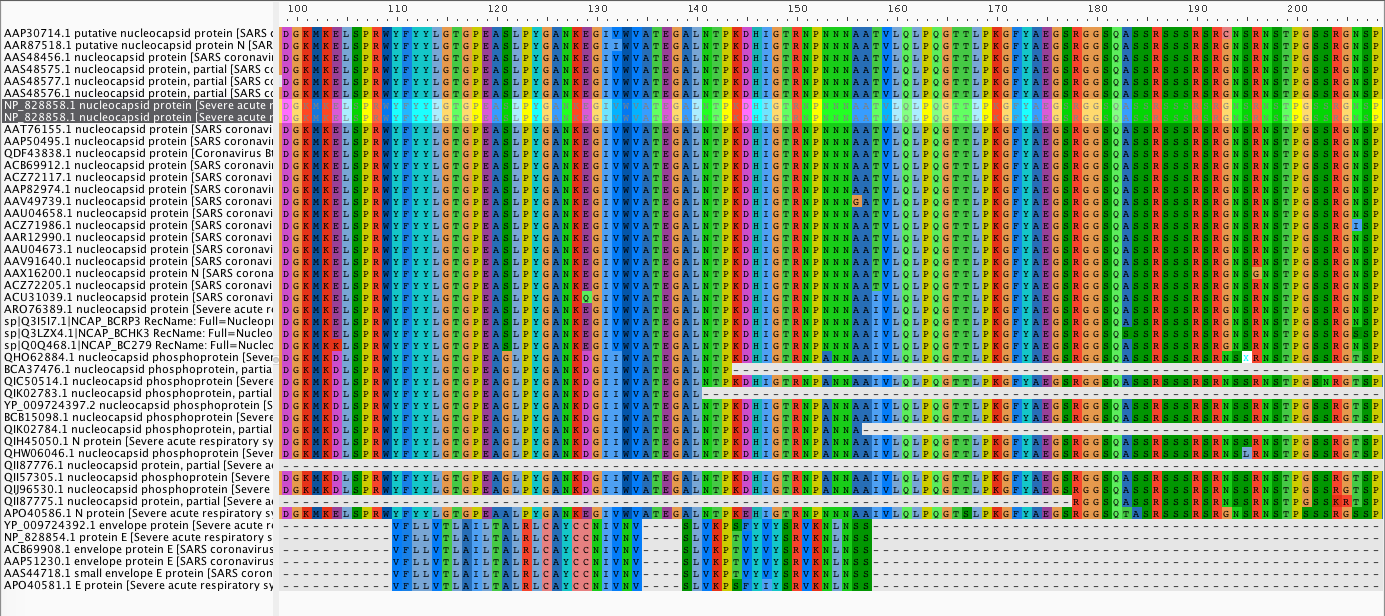

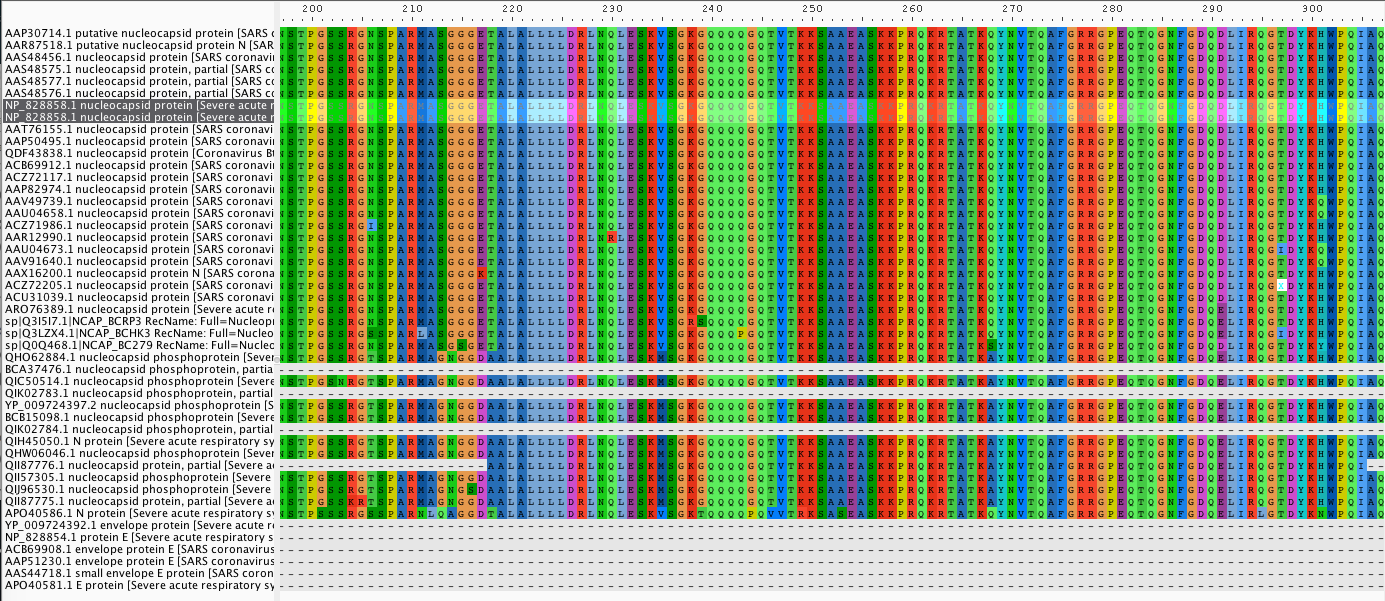

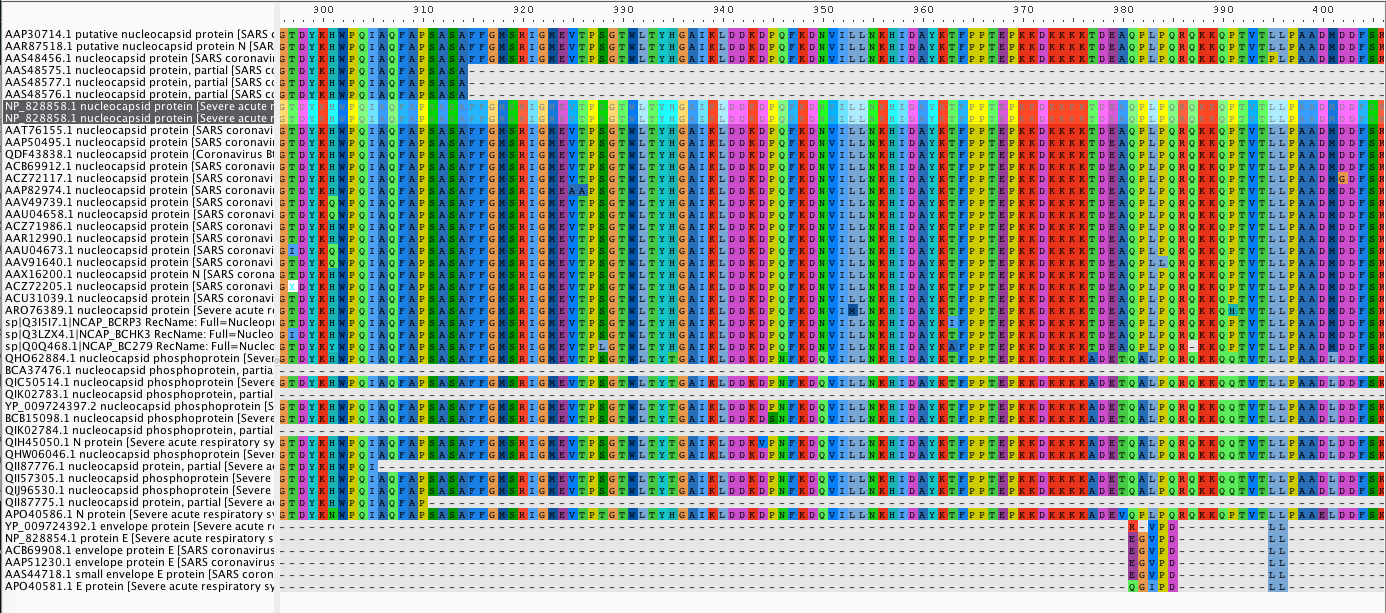

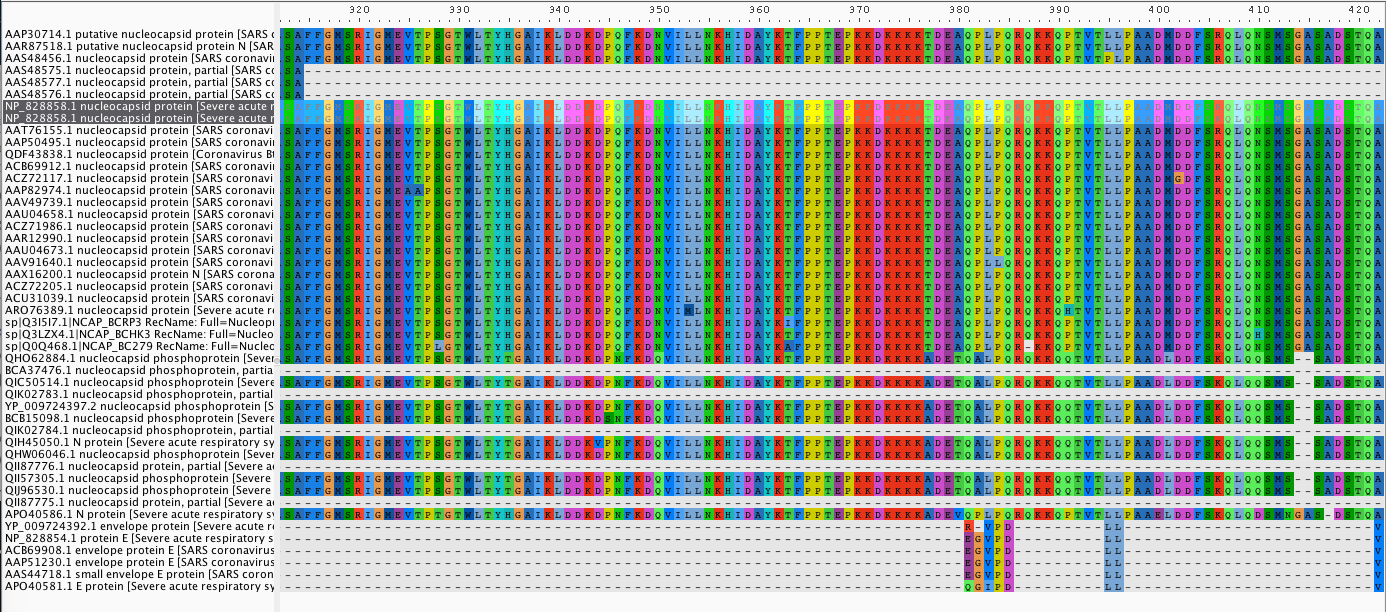

Removed sequences:

NP_828858.1 nucleocapsid protein [Severe acute respiratory syndrome-related coronavirus] – duplicate

AAS48577.1 nucleocapsid protein, partial [SARS coronavirus cw037] – partial

AAS48575.1 nucleocapsid protein, partial [SARS coronavirus xw002] – partial

AAS48576.1 nucleocapsid protein, partial [SARS coronavirus cw049] – partial

BCA37476.1 nucleocapsid phosphoprotein, partial [Severe acute respiratory syndrome coronavirus 2] – partial

QII87776.1 nucleocapsid protein, partial [Severe acute respiratory syndrome coronavirus 2] – partial

QII87775.1 nucleocapsid protein, partial [Severe acute respiratory syndrome coronavirus 2] – partial

YP_009724392.1 envelope protein [Severe acute respiratory syndrome coronavirus 2] – partial

NP_828854.1 protein E [Severe acute respiratory syndrome-related coronavirus] – partial

ACB69908.1 envelope protein E [SARS coronavirus BJ182-12] – partial

AAP51230.1 envelope protein E [SARS coronavirus GD01] – partial

AAS44718.1 small envelope E protein [SARS coronavirus TW-GD1] – partial

APO40581.1 E protein [Severe acute respiratory syndrome-related coronavirus] – partial

QIK02783.1 nucleocapsid phosphoprotein, partial [Severe acute respiratory syndrome coronavirus 2] – partial

QIK02784.1 nucleocapsid phosphoprotein, partial [Severe acute respiratory syndrome coronavirus 2] – partial

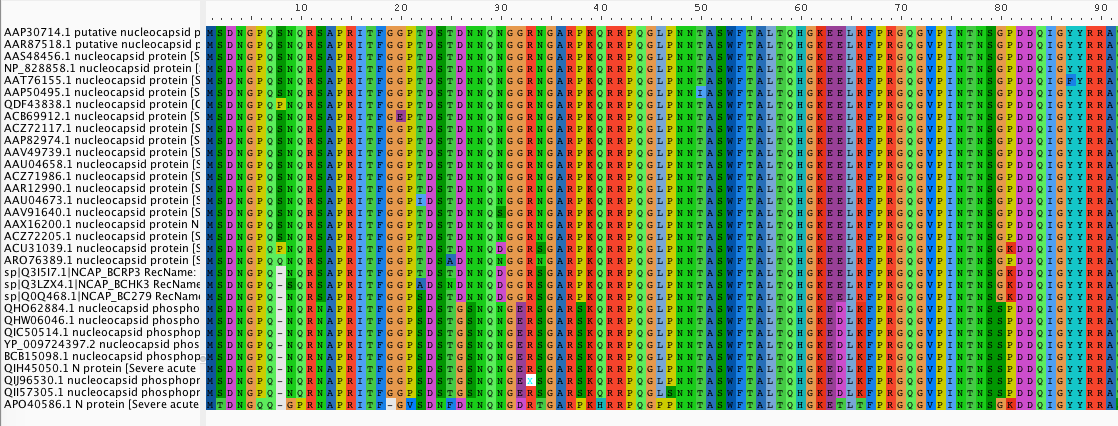

***Part 3. Alignment of Spike Protein Variants***

March 30, 2020

Spike Protein – SARS CoV-2

Alignment with MAFFT and visualized with AliView

The sequences used in this alignment:

>sp|P59594.1|SPIKE_CVHSA RecName: Full=Spike glycoprotein; Short=S glycoprotein; AltName: Full=E2; AltName: Full=Peplomer protein; Contains: RecName: Full=Spike protein S1; Contains: RecName: Full=Spike protein S2; Contains: RecName: Full=Spike protein S2'; Flags: Precursor

MFIFLLFLTLTSGSDLDRCTTFDDVQAPNYTQHTSSMRGVYYPDEIFRSDTLYLTQDLFLPFYSNVTGFH

TINHTFGNPVIPFKDGIYFAATEKSNVVRGWVFGSTMNNKSQSVIIINNSTNVVIRACNFELCDNPFFAV

SKPMGTQTHTMIFDNAFNCTFEYISDAFSLDVSEKSGNFKHLREFVFKNKDGFLYVYKGYQPIDVVRDLP

SGFNTLKPIFKLPLGINITNFRAILTAFSPAQDIWGTSAAAYFVGYLKPTTFMLKYDENGTITDAVDCSQ

NPLAELKCSVKSFEIDKGIYQTSNFRVVPSGDVVRFPNITNLCPFGEVFNATKFPSVYAWERKKISNCVA

DYSVLYNSTFFSTFKCYGVSATKLNDLCFSNVYADSFVVKGDDVRQIAPGQTGVIADYNYKLPDDFMGCV

LAWNTRNIDATSTGNYNYKYRYLRHGKLRPFERDISNVPFSPDGKPCTPPALNCYWPLNDYGFYTTTGIG

YQPYRVVVLSFELLNAPATVCGPKLSTDLIKNQCVNFNFNGLTGTGVLTPSSKRFQPFQQFGRDVSDFTD

SVRDPKTSEILDISPCSFGGVSVITPGTNASSEVAVLYQDVNCTDVSTAIHADQLTPAWRIYSTGNNVFQ

TQAGCLIGAEHVDTSYECDIPIGAGICASYHTVSLLRSTSQKSIVAYTMSLGADSSIAYSNNTIAIPTNF

SISITTEVMPVSMAKTSVDCNMYICGDSTECANLLLQYGSFCTQLNRALSGIAAEQDRNTREVFAQVKQM

YKTPTLKYFGGFNFSQILPDPLKPTKRSFIEDLLFNKVTLADAGFMKQYGECLGDINARDLICAQKFNGL

TVLPPLLTDDMIAAYTAALVSGTATAGWTFGAGAALQIPFAMQMAYRFNGIGVTQNVLYENQKQIANQFN

KAISQIQESLTTTSTALGKLQDVVNQNAQALNTLVKQLSSNFGAISSVLNDILSRLDKVEAEVQIDRLIT

GRLQSLQTYVTQQLIRAAEIRASANLAATKMSECVLGQSKRVDFCGKGYHLMSFPQAAPHGVVFLHVTYV

PSQERNFTTAPAICHEGKAYFPREGVFVFNGTSWFITQRNFFSPQIITTDNTFVSGNCDVVIGIINNTVY

DPLQPELDSFKEELDKYFKNHTSPDVDLGDISGINASVVNIQKEIDRLNEVAKNLNESLIDLQELGKYEQ

YIKWPWYVWLGFIAGLIAIVMVTILLCCMTSCCSCLKGACSCGSCCKFDEDDSEPVLKGVKLHYT

>ACZ72035.1 spike glycoprotein precursor [SARS coronavirus MA15]

MFIFLLFLTLTSGSDLDRCTTFDDVQAPNYTQHTSSMRGVYYPDEIFRSDTLYLTQDLFLPFYSNVTGFH

TINHTFGNPVIPFKDGIYFAATEKSNVVRGWVFGSTMNNKSQSVIIINNSTNVVIRACNFELCDNPFFAV

SKPMGTQTHTMIFDNAFNCTFEYISDAFSLDVSEKSGNFKHLREFVFKNKDGFLYVYKGYQPIDVVRDLP

SGFNTLKPIFKLPLGINITNFRAILTAFSPAQDIWGTSAAAYFVGYLKPTTFMLKYDENGTITDAVDCSQ

NPLAELKCSVKSFEIDKGIYQTSNFRVVPSGDVVRFPNITNLCPFGEVFNATKFPSVYAWERKKISNCVA

DYSVLYNSTFFSTFKCYGVSATKLNDLCFSNVYADSFVVKGDDVRQIAPGQTGVIADYNYKLPDDFMGCV

LAWNTRNIDATSTGNHNYKYRYLRHGKLRPFERDISNVPFSPDGKPCTPPALNCYWPLNDYGFYTTTGIG

YQPYRVVVLSFELLNAPATVCGPKLSTDLIKNQCVNFNFNGLTGTGVLTPSSKRFQPFQQFGRDVSDFTD

SVRDPKTSEILDISPCSFGGVSVITPGTNASSEVAVLYQDVNCTDVSTAIHADQLTPAWRIYSTGNNVFQ

TQAGCLIGAEHVDTSYECDIPIGAGICASYHTVSLLRSTSQKSIVAYTMSLGADSSIAYSNNTIAIPTNF

SISITTEVMPVSMAKTSVDCNMYICGDSTECANLLLQYGSFCTQLNRALSGIAAEQDRNTREVFAQVKQM

YKTPTLKYFGGFNFSQILPDPLKPTKRSFIEDLLFNKVTLADAGFMKQYGECLGDINARDLICAQKFNGL

TVLPPLLTDDMIAAYTAALVSGTATAGWTFGAGAALQIPFAMQMAYRFNGIGVTQNVLYENQKQIANQFN

KAISQIQESLTTTSTALGKLQDVVNQNAQALNTLVKQLSSNFGAISSVLNDILSRLDKVEAEVQIDRLIT

GRLQSLQTYVTQQLIRAAEIRASANLAATKMSECVLGQSKRVDFCGKGYHLMSFPQAAPHGVVFLHVTYV

PSQERNFTTAPAICHEGKAYFPREGVFVFNGTSWFITQRNFFSPQIITTDNTFVSGNCDVVIGIINNTVY

DPLQPELDSFKEELDKYFKNHTSPDVDLGDISGINASVVNIQKEIDRLNEVAKNLNESLIDLQELGKYEQ

YIKWPWYVWLGFIAGLIAIVMVTILLCCMTSCCSCLKGACSCGSCCKFDEDDSEPVLKGVKLHYT

>AAP50485.1 spike glycoprotein [SARS coronavirus FRA]

MFIFLLFLTLTSGSDLDRCTTFDDVQAPNYTQHTSSMRGVYYPDEIFRSDTLYLTQDLFLPFYSNVTGFH

TINHTFGNPVIPFKDGIYFAATEKSNVVRGWVFGSTMNNKSQSVIIINNSTNVVIRACNFELCDNPFFAV

SKPMGTQTHTMIFDNAFNCTFEYISDAFSLDVSEKSGNFKHLREFVFKNKDGFLYVYKGYQPIDVVRDLP

SGFNTLKPIFKLPLGINITNFRAILTAFSPAQDIWGTSAAAYFVGYLKPTTFMLKYDENGTITDAVDCSQ

NPLAELKCSVKSFEIDKGIYQTSNFRVVPSGDVVRFPNITNLCPFGEVFNATKFPSVYAWERKKISNCVA

DYSVLYNSTFFSTFKCYGVSATKLNDLCFSNVYADSFVVKGDDVRQIAPGQTGVIADYNYKLPDDFMGCV

LAWNTRNIDATSTGNYNYKYRYLRHGKLRPFERDISNVPFSPDGKPCTPPALNCYWPLNDYGFYTTTGIG

YQPYRVVVLSFELLNAPATVCGPKLSTDLIKNQCVNFNFNGLTGTGVLTPSSKRFQPFQQFGRDVSDFTD

SVRDPKTSEILDISPCSFGGVSVITPGTNASSEVAVLYQDVNCTDVSTAIHADQLTPAWRIYSTGNNVFQ

TQAGCLIGAEHVDTSYECDIPIGAGICASYHTVSLLRSTSQKSIVAYTMSLGADSSIAYSNNTIAIPTNF

SISITTEVMPVSMAKTSVDCNMYICGDSTECANLLLQYGSFCTQLNRALSGIAAEQDRNTREVFAQVKQM

YKTPTLKYFGGFNFSQILPDPLKPTKRSFIEDLLFNKVTLADAGFMKQYGECLGDINARDLICAQKFNGL

TVLPPLLTDDMIAAYTAALVSGTATAGWTFGAGAALQIPFAMQMAYRFNGIGVTQNVLYENQKQIANQFN

KAISQIQESLTTTSTALGKLQDVVNQNAQALNTLVKQLSSNFGAISSVLNDILSRLDKVEAEVQIDRLIT

GRLQSLQTYVTQQLIRAAEIRASANLAATKMSECVLGQSKRVDFCGKGYHLMSFPQAAPHGVVFLHVTYV

PSQERNFTTAPAICHEGKAYFPREGVFVFNGTSWFITQRNFFSPQIITTDNTFVSGNCDVVIGIINNTVY

DPLQPELDSFKEELDKYFKNHTSPDVDFGDISGINASVVNIQKEIDRLNEVAKNLNESLIDLQELGKYEQ

YIKWPWYVWLGFIAGLIAIVMVTILLCCMTSCCSCLKGACSCGSCCKFDEDDSEPVLKGVKLHYT

>AAP30030.1 spike glycoprotein S [SARS coronavirus BJ01]

MFIFLLFLTLTSGSDLDRCTTFDDVQAPNYTQHTSSMRGVYYPDEIFRSDTLYLTQDLFLPFYSNVTGFH

TINHTFDNPVIPFKDGIYFAATEKSNVVRGWVFGSTMNNKSQSVIIINNSTNVVIRACNFELCDNPFFAV

SKPMGTQTHTMIFDNAFNCTFEYISDAFSLDVSEKSGNFKHLREFVFKNKDGFLYVYKGYQPIDVVRDLP

SGFNTLKPIFKLPLGINITNFRAILTAFSPAQDTWGTSAAAYFVGYLKPTTFMLKYDENGTITDAVDCSQ

NPLAELKCSVKSFEIDKGIYQTSNFRVVPSGDVVRFPNITNLCPFGEVFNATKFPSVYAWERKKISNCVA

DYSVLYNSTFFSTFKCYGVSATKLNDLCFSNVYADSFVVKGDDVRQIAPGQTGVIADYNYKLPDDFMGCV

LAWNTRNIDATSTGNYNYKYRYLRHGKLRPFERDISNVPFSPDGKPCTPPALNCYWPLNDYGFYTTTGIG

YQPYRVVVLSFELLNAPATVCGPKLSTDLIKNQCVNFNFNGLTGTGVLTPSSKRFQPFQQFGRDVSDFTD

SVRDPKTSEILDISPCSFGGVSVITPGTNASSEVAVLYQDVNCTDVSTAIHADQLTPAWRIYSTGNNVFQ

TQAGCLIGAEHVDTSYECDIPIGAGICASYHTVSLLRSTSQKSIVAYTMSLGADSSIAYSNNTIAIPTNF

SISITTEVMPVSMAKTSVDCNMYICGDSTECANLLLQYGSFCTQLNRALSGIAAEQDRNTREVFAQVKQM

YKTPTLKYFGGFNFSQILPDPLKPTKRSFIEDLLFNKVTLADAGFMKQYGECLGDINARDLICAQKFNGL

TVLPPLLTDDMIAAYTAALVSGTATAGWTFGAGAALQIPFAMQMAYRFNGIGVTQNVLYENQKQIANQFN

KAISQIQESLTTTSTALGKLQDVVNQNAQALNTLVKQLSSNFGAISSVLNDILSRLDKVEAEVQIDRLIT

GRLQSLQTYVTQQLIRAAEIRASANLAATKMSECVLGQSKRVDFCGKGYHLMSFPQAAPHGVVFLHVTYV

PSQERNFTTAPAICHEGKAYFPREGVFVFNGTSWFITQRNFFSPQIITTDNTFVSGNCDVVIGIINNTVY

DPLQPELDSFKEELDKYFKNHTSPDVDLGDISGINASVVNIQKEIDRLNEVAKNLNESLIDLQELGKYEQ

YIKWPWYVWLGFIAGLIAIVMVTILLCCMTSCCSCLKGACSCGSCCKFDEDDSEPVLKGVKLHYT

>AAP51227.1 spike glycoprotein S [SARS coronavirus GD01]

MFIFLLFLTLTSGSDLDRCTTFDDVQAPNYTQHTSSMRGVYYPDEIFRSDTLYLTQDLFLPFYSNVTGFH

TINHTFDNPVIPFKDGIYFAATEKSNVVRGWVFGSTMNNKSQSVIIINNSTNVVIRACNFELCDNPFFAV

SKPMGTQTHTMIFDNAFNCTFEYISDAFSLDVSEKSGNFKHLREFVFKNKDGFLYVYKGYQPIDVVRDLP

SGFNTLKPIFKLPLGINITNFRAILTAFLPAQDTWGTSAAAYFVGYLKPTTFMLKYDENGTITDAVDCSQ

NPLAELKCSVKSFEIDKGIYQTSNFRVVPSRDVVRFPNITNLCPFGEVFNATKFPSVYAWERKRISNCVA

DYSVLYNSTFFSTFKCYGVSATKLNDLCFSNVYADSFVVKGDDVRQIAPGQTGVIADYNYKLPDDFMGCV

LAWNTRNIDATSTGNYNYKYRYLRHGKLRPFERDISNVPFSPDGKPCTPPALNCYWPLNDYGFYTTTGIG

YQPYRVVVLSYELLNAPATVCGPKLSTDLIKNQCVNFNFNGLTGTGVLTPSSKRFQPFQQFGRDVSDFTD

SVRDPKTSEILDISPCSFGGVSVITPGTNASSEVAVLYQDVNCTDVSTAIHADQLTPAWRIYSTGNNVFQ

TQAGCLIGAEHVDTSYECDIPIGAGICASYHTVSLLRSTSQKSIVAYTMSLGADSSIAYSNNTIAIPTNF

SISITTEVMPVSMAKTSVDCNMYICGDSTECANLLLQYGSFCTQLNRALSGIAAEQDRNTREVFAQVKQM

YKTPTLKDFGGFNFSQILPDPLKSTKRSFIEDLLFNKVTLADAGFMKQYGECLGDINARDLICAQKFNGL

TVLPPLLTDDMIAAYTAALVSGTATAGWTFGAGAALQIPFAMQMAYRFNGIGVTQNVLYENQKQIANQFN

KAISQIQESLTTTSTALGKLQDVVNQNAQALNTLVKQLSSNFGAISSVLNDILSRLDKVEAEVQIDRLIT

GRLQSLQTYVTQQLIRAAEIRASANLAATKMSECVLGQSKRVDFCGKGYHLMSFPQAAPHGVVFLHVTYV

PSQERNFTTAPAICHEGKAYFPREGVFVFNGTSWFITQRNFFSPQIITTDNTFVSGNCDVVIGIINNTVY

DPLQPELDSFKEELDKYFKNHTSPDVDLGDISGINASVVNIQKEIDRLNEVAKNLNESLIDLQELGKYEQ

YIKWPWYVWLGFIAGLIAIVMVTILLCCMTSCCSCLKGACSCGSCCKFDEDDSEPVLKGVKLHYT

>AAX16192.1 spike glycoprotein S [SARS coronavirus WH20]

MFIFLLFLTLTSGSDLDRCTTFDDVQAPNYTQHTSSMRGVYYPDEIFRSDTLYLTQDLFLPFYSNVTGFH

TINHTFDNPVIPFKDGIYFAATEKSNVVRGWVFGSTMNNKSQSVIIINNSTNVVIRACNFELCDNPFFAV

SKPMGTQTHTMIFDNAFNCTFEYISDAFSLDVSEKSGNFKHLREFVFKNKDGFLYVYKGYQPIDVVRDLP

SGFNTLKPIFKLPLGINITNFRAILTAFSPAQDTWGTSAAAYFVGYLKPTTFMLKYDENGTITDAVDCSQ

NPLAELKCSVKSFEIDKGIYQTSNFRVVPSGDVVRFPNITNLCPFGEVFNATKFPSVYAWERKKISNCVA

DYSVLYNSTFFSTFKCYGVSATKLNDLCFSNVYADSFVVKGDDVRQIAPGQTGVIADYNYKLPDDFMGCV

LAWNTRNIDATSTGNYNYKYRYLRHGKLRPFERDISNVPFSPDGKPCTPPALNCYWPLNDYGFYTTTGIG

YQPYRVVVLSFELLNAPATVCGPKLSTDLIKNQCVNFNFNGLTGTGVLTPSSKRFQPFQQFGRDVSDFTD

SVRDPKTSEILDISPCSFGGVSVITPGTNASSEVAVLYQDVNCTDVSTAIHADQLTPAWRIYSTGNNVFQ

TQAGCLIGAEHVDTSYECDIPIGAGICASYHTVSLLRSTSQKSIVAYTMSLGADSSIAYSNNTIAIPTNF

SISITTEVMPVSMAKTSVDCNMYICGDSTECANLLLQYGSFCTQLNRALSGIAAEQDRNTREVFAQVKQM

YKTPTLKYFGGFNFSQILPDPLKPTKRSFIEDLLFNKVTLADAGFMKQYGECLGDINARDLICAQKFNGL

TVLPPLLTDDMIAAYTAALVSGTATAGWTFGAGAALQIPFAMQMAYRFNGIGVTQNVLYENQKQIANQFN

KAISQIQESLTTTSTALGKLQDVVNQNAQALNTLVKQLSSNFGAISSVLNDILSRLDKVEAEVQIDRLIT

GRLQSLQTYVTQQLIRAAEIRASANLAATKMSECVLGQSKRVDFCGKGYHLMSFPQAAPHGVVFLHVTYV

PSQERNFTTAPAICHEGKAYFPREGVFVFNGTSWFITQRNFFSPQIITTDNTFVSGNCDVVIGIINNTVY

DPLQPELDSFKEELDKYFKNHTSPDVDLGDISGINASVVNIQKEIDRLNEVAKNLNESLIDLQELGKYEQ

YIKWPWYVWLGFIAGLMAIVMVTILLCCMTSCCSCLKGACSCGSCCKFDEDDSEPVLKGVKLHYT

>ACB69905.1 spike glycoprotein S [SARS coronavirus BJ182-12]

MFIFLLFLTLTSGSDLDRCTTFDDVQAPNYTQHTSSMRGVYYPDEIFRSDTLYLTQDLFLPFYSNVTGFH

TINHTFGNPVIPFKDGIYFAATEKSNVVRGWVFGSTMNNKSQSVIIINNSTNVVIRACNFELCDNPFFAV

SKPMGTQTHTMIFDNAFNCTFEYISDAFSLDVSEKSGNFKHLREFVFKNKDGFLYVYKGYQPIDVVRDLP

SGFNTLKPIFKLPLGINITNFRAILTAFSPAQDIWGTSAAAYFVGYLKPTTFMLKYDENGTITDAVDCSQ

NPLAELKCSVKSFEIDKGIYQTSNFRVVPSGDVVRFPNITNLCPFGEVFNATKFPSVYAWERKKISNCVA

DYSVLYNSTFFSTFKCYGVSATKLNDLCFSNVYADSFVVKGDDVRQIAPGQTGVIADYNYKLPDDFMGCV

LAWNTRNIDATSTGNYNYKYRYLRHGKLRPFERDISNVPFSPDGKPCTPPALNCYWPLNDYGFYTTTGIG

YQPYRVVVLSFELLNAPATVCGPKLSTDLIKNQCVNFNFNGLTGTGVLTPSSKRFQPFQQFGRDVSDFTD

SVRDPKTSEILDISPCSFGGVSVITPGTNASSEVAVLYQDVNCTDVSTAIHADQLTPAWRIYSTGNNVFQ

TQAGCLIGAEHVDTSYECDIPIGAGICASYHTVSLLRSTSQKSIVAYTMSLGADSSIAYSNNTIAIPTNF

SISITTEVMPVSMAKTSVDCNMYICGDSTECANLLLQYGSFCTQLNRALSGIAAEQDRNTREVFAQVKQM

YKTPTLKYFDGFNFSQILPDPLKPIKRSFIEDLLFNKVTLADAGFMKQYGECLGDINARDLICAQKFNGL

TVLPPLLTDDMIAAYTAALVSGTATAGWTFGAGAALQIPFAMQMAYRFNGIGVTQNVLYENQKQIANQFN

KAISQIQESLTTTSTALGKLQDVVNQNAQALNTLVKQLSSNFGAISSVLNDILSRLDKVEAEVQIDRLIT

GRLQSLQTYVTQQLIRAAEIRASANLAATKMSECVLGQSKRVDFCGKGYHLMSFPQAAPHGVVFLHVTYV

PSQERNFTTAPAICHEGKAYFPREGVFVFNGTSWFITQRNFFSPQIITTDNTFVSGNCDVVIGIINNTVY

DPLQPELDSFKEELDKYFKNHTSPDVDLGDISGINAPVVNIQKEIDRLNEVAKNLNESLIDLRELGKYEQ

YIKWPWYVWLGFIAGLIAIVMVTILLCCMTSCCSCLKGACSCGSCCKFDEDDSEPVLKGVKLHYT

>ACB69894.1 spike glycoprotein S [SARS coronavirus BJ182-8]

MFIFLLFLTLTSGSDLDRCTTFDDVQAPNYTQHTSSMRGVYYPDEIFRSDTLYLTQDLFLPFYSNVTGFH

TINHTFGNPVIPFKDGIYFAATEKSNVVRGWVFGSTMNNKSQSVIIINNSTNVVIRACNFELCDNPFFAV

SKPMGTQTHTMIFDNAFNCTFEYISDAFSLDVSEKSGNFKHLREFVFKNKDGFLYVYKGYQPIDVVRDLP

SGFNTLKPIFKLPLGINITNFRAILTAFSPAQDIWGTSAAAYFVGYLKPTTFMLKYDENGTITDAVDCSQ

NPLAELKCSVKSFEIDKGIYQTSNFRVVPSGDVVRFPNITNLCPFGEVFNATKFPSVYAWERKKISNCVA

DYSVLYNSTFFSTFKCYGVSATKLNDLCFSNVYADSFVVKGDDVRQIAPGQTGVIADYNYKLPDDFMGCV

LAWNTRNIDATSTGNYNYKYRYLRHGKLRPFERDISNVPFSPDGKPCTPPALNCYWPLNDYGFYTTTGIG

YQPYRVVVLSFELLNAPATVCGPKLSTDLIKNQCVNFNFNGLTGTGVLTPSSKRFQPFQQFGRDVSDFTD

SVRDPKTSEILDISPCSFGGVSVITPGTNASSEVAVLYQDVNCTDVSTAIHADQLTPAWRIYSTGNNVFQ

TQAGCLIGAEHVDTSYECDIPIGAGICASYHTVSLLRSTSQKSIVAYTMSLGADSSIAYSNNTIAIPTNF

SISITTEVMPVSMAKTSVDCNMYICGDSTECANLLLQYGSFCTQLNRALSGIAAEQDRNTREVFAQVKQM

YKTPTLKYFDGFNFSQILPDPLKPTKRSFIEDLLFNKVTLADAGFMKQYGECLGDINARDLICAQKFNGL

TVLPPLLTDDMIAAYTAALVSGTATAGWTFGAGAALQIPFAMQMAYRFNGIGVTQNVLYENQKQIANQFN

KAISQIQESLTTTSTALGKLQDVVNQNAQALNTLVKQLSSNFGAISSVLNDILSRLDKVEAEVQIDRLIT

GRLQSLQTYVTQQLIRAAEIRASANLAATKMSECVLGQSKRVDFCGKGYHLMSFPQAAPHGVVFLHVTYV

PSQERNFTTAPAICHEGKAYFPREGVFVFNGTSWFITQRNFFSPQIITTDNTFVSGNCDVVIGIINNTVY

DPLQPELDSFKEELDKYFKNHTSPDVDLGDISGINASVVNIQKEIDRLNEVAKNLNESLIDLRELGKYEQ

YIKWPWYVWLGFIAGLIAIVMVTILLCCMTSCCSCLKGACSCGSCCKFDEDDSEPVLKGVKLHYT

>ACB69883.1 spike glycoprotein S [SARS coronavirus BJ182-4]

MFIFLLFLTLTSGSDLDRCTTFDDVQAPNYTQHTSSMRGVYYPDEIFRSDTLYLTQDLFLPFYSNVTGFH

TINHTLGNPVIPFKDGIYFAATEKSNVVRGWVFGSTMNNKSQSVIIINNSTNVVIRACNFELCDNPFFAV

SKPMGTQTHTMIFDNAFNCTFEYISDAFSLDVSEKSGNFKHLREFVFKNKDGFLYVYKGYQPIDVVRDLP

SGFNTLKPIFKLPLGINITNFRAILTAFSPAQDIWGTSAAAYFVGYLKPTTFMLKYDENGTITDAVDCSQ

NPLAELKCSVKSFEIDKGIYQTSNFRVVPSGDVVRFPNITNLCPFGEVFNATKFPSVYAWERKKISNCVA

DYSVLYNSTFFSTFKCYGVSATKLNDLCFSNVYADSFVVKGDDVRQIAPGQTGVIADYNYKLPDDFMGCV

LAWNTRNIDATSTGNYNYKYRYLRHGKLRPFERDISNVPFSPDGKPCTPPALNCYWPLNDYGFYTTTGIG

YQPYRVVVLSFELLNAPATVCGPKLSTDLIKNQCVNFNFNGLTGTGVLTPSSKRFQPFQQFGRDVSDFTD

SVRDPKTSEILDISPCSFGGVSVITPGTNASSEVAVLYQDVNCTDVSTAIHADQLTPAWRIYSTGNNVFQ

TQAGCLIGAEHVDTSYECDIPIGAGICASYHTVSLLRSTSQKSIVAYTMSLGADSSIAYSNNTIAIPTNF

SISITTEVMPVSMAKTSVDCNMYICGDSTECANLLLQYGSFCTQLNRALSGIAAEQDRNTREVFAQVKQM

YKTPTLKYFDGFNFSQILPDPLKPTKRSFIEDLLFNKVTLADAGFMKQYGECLGDINARDLICAQKFNGL

TVLPPLLTDDMIAAYTAALVSGTATAGWTFGAGAALQIPFAMQMAYRFNGIGVTQNVLYENQKQIANQFN

KAISQIQESLTTTSTALGKLQDVVNQNAQALNTLVKQLSSNFGAISSVLNDILSRLDKVEAEVQIDRLIT

GRLQSLQTYVTQQLIRAAEIRASANLAATKMSECVLGQSKRVDFCGKGYHLMSFPQAAPHGVVFLHVTYV

PSQERNFTTAPAICHEGKAYFPREGVFVFNGTSWFITQRNFFSPQIITTDNTFVSGNCDVVIGIINNTVY

DPLQPELDSFKEELDKYFKNHTSPDVDLGDISGINASVVNIQKEIDRLNEVAKNLNESLIDLRELGKYEQ

YIKWPWYVWLGFIAGLIAIVMVTILLCCMTSCCSCLKGACSCGSCCKFDEDDSEPVLKGVKLHYT

>ACB69860.1 spike glycoprotein S [SARS coronavirus BJ182a]

MFIFLLFLTLTSGSDLDRCTTFDDVQAPNYTQHTSSMRGVYYPDEIFRSDTLYLTQDLFLPFYSNVTGFH

TINHTFGNPIIPFKDGIYFAATEKSNVVRGWVFGSTMNNKSQSVIIINNSTNVVIRACNFELCDNPFFAV

SKPMGTQTHTMIFDNAFNCTFEYISDAFSLDVSEKSGNFKHLREFVFKNKDGFLYVYKGYQPIDVVRDLP

SGFNTLKPIFKLPLGINITNFRAILTAFSPAQDIWGTSAAAYFVGYLKPTTFMLKYDENGTITDAVDCSQ

NPLAELKCSVKSFEIDKGIYQTSNFRVVPSGDVVRFPNITNLCPFGEVFNATKFPSVYAWERKKISNCVA

DYSVLYNSTFFSTFKCYGVSATKLNDLCFSNVYADSFVVKGDDVRQIAPGQTGVIADYNYKLPDDFMGCV

LAWNTRNIDATSTGNYNYKYRYLRHGKLRPFERDISNVPFSPDGKPCTPPALNCYWPLNDYGFYTTTGIG

YQPYRVVVLSFELLNAPATVCGPKLSTDLIKNQCVNFNFNGLTGTGVLTPSSKRFQPFQQFGRDVSDFTD

SVRDPKTSEILDISPCSFGGVSVITPGTNASSEVAVLYQDVNCTDVSTAIHADQLTPAWRIYSTGNNVFQ

TQAGCLIGAEHVDTSYECDIPIGAGICASYHTVSLLRSTSQKSIVAYTMSLGADSSIAYSNNTIAIPTNF

SISITTEVMPVSMAKTSVDCNMYICGDSTECANLLLQYGSFCTQLNRALSGIAAEQDRNTREVFAQVKQM

YKTPTLKYFDGFNFSQILPDPLKPTKRSFIEDLLFNKVTLADAGFMKQYGECLGDINARDLICAQKFNGL

TVLPPLLTDDMIAAYTAALVSGTATAGWAFGAGAALQIPFAMQMAYRFNGIGVTQNVLYENQKQIANQFN

KAISQIQESLTTTSTALGKLQDVVNQNAQALNTLVKQLSSNFGAISSVLNDILSRLDKVEAEVQIDRLIT

GRLQSLQTYVTQQLIRAAEIRASANLAATKMSECVLGQSKRVDFCGKGYHLMSFPQAAPHGVVFLHVTYV

PSQERNFTTAPAICHEGKAYFPREGVFVFNGTSWFITQRNFFSPQIITTDNTFVSGNCDVVIGIINNTVY

DPLQPELDSFKEELDKYFKNHTSPDVDLGDISGINASVVNIQKEIDRLNEVAKNLNESLIDLRELGKYEQ

YIKWPWYVWLGFIAGLIAIVMVTILLCCMTSCCSCLKGACSCGSCCKFDEDDSEPVLKGVKLHYT

>AAU81608.1 S protein [SARS Coronavirus CDC#200301157]

MFIFLLFLTLTSGSDLDRCTTFDDVQAPNYTQHTSSMRGVYYPDEIFRSDTLYLTQDLFLPFYSNVTGFH

TINHTFGNPVIPFKDGIYFAATEKSNVVRGWVFGSTMNNKSQSVIIINNSTNVVIRACNFELCDNPFFAV

SKPMGTQTHTMIFDNAFNCTFEYISDAFSLDVSEKSGNFKHLREFVFKNKDGFLYVYKGYQPIDVVRDLP

SGFNTLKPIFKLPLGINITNFRAILTAFSPAQDIWGTSAAAYFVGYLKPTTFMLKYDENGTITDAVDCSQ

NPLAELKCSVKSFEIDKGIYQTSNFRVVPSGDVVRFPNITNLCPFGEVFNATKFPSVYAWERKKISNCVA

DYSVLYNSTFFSTFKCYGVSATKLNDLCFSNVYADSFVVKGDDVRQIAPGQTGVIADYNYKLPDDFMGCV

LAWNTRNIDATSTGNYNYKYRYLRHGKLRPFERDISNVPFSPDGKPCTPPALNCYWPLNDYGFYTTTGIG

YQPYRVVVLSFELLNAPATVCGPKLSTDLIKNQCVNFNFNGLTGTGVLTPSSKRFQPFQQFGRDVSDFTD

SVRDPKTSEILDISPCSFGGVSVITPGTNASSEVAVLYQDVNCTDVSTAIHADQLTPAWRIYSTGNNVFQ

TQAGCLIGAEHVDTSYECDIPVGAGICASYHTVSLLRSTSQKSIVAYTMSLGADSSIAYSNNTIAIPTNF

SISITTEVMPVSMAKTSVDCNMYICGDSTECANLLLQYGSFCTQLNRALSGIAAEQDRNTREVFAQVKQM

YKTPTLKYFGGFNFSQILPDPLKPTKRSFIEDLLFNKVTLADAGFMKQYGECLGDINARDLICAQKFNGL

TVLPPLLTDDMIAAYTAALVSGTATAGWTFGAGAALQIPFAMQMAYRFNGIGVTQNVLYENQKQIANQFN

KAISQIQESLTTTSTALGKLQDVVNQNAQALNTLVKQLSSNFGAISSVLNDILSRLDKVEAEVQIDRLIT

GRLQSLQTYVTQQLIRAAEIRASANLAATKMSECVLGQSKRVDFCGKGYHLMSFPQAAPHGVVFLHVTYV

PSQERNFTTAPAICHEGKAYFPREGVFVFNGTSWFITQRNFFSPQIITTDNTFVSGNCDVVIGIINNTVY

DPLQPELDSFKEELDKYFKNHTSPDVDLGDISGINASVVNIQKEIDRLNEVAKNLNESLIDLQELGKYEQ

YIKWPWYVWLGFIAGLIAIVMVTILLCCMTSCCSCLKGACSCGSCCKFDEDDSEPVLKGVKLHYT

>AAR91586.1 spike glycoprotein S [SARS coronavirus NS-1]

MFIFLLFLTLTSGSDLDRCTTFDDVQAPNYTQHTSSMRGVYYPDEIFRSDTLYLTQDLFLPFYSNVTGFH

TINHTFGNPVIPFKDGIYFAATEKSNVVRGWVFGSTMNNKSQSVIIINNSTNVVIRACNFELCDNPFFAV

SKPMGTQTHTMIFDNAFNCTFEYISDAFSLDVSEKSGNFKHLREFVFKNKDGFLYVYKGYQPIDVVRDLP

SGFNTLKPIFKLPLGINITNFRAILTAFSPAQDTWGTSAAAYFVGYLKPTTFMLKYDENGTITDAVDCSQ

NPLAELKCSVKSFEIDKGIYQTSNFRVVPSGDVVRFPNITNLCPFGEVFNATKFPSVYAWERKKISNCVA

DYSVLYNSTFFSTFKCYGVSATKLNDLCFSNVYADSFVVKGDDVRQIAPGQTGVIADYNYKLPDDFMGCV

LAWNTRNIDATSTGNYNYKYRYLRHGKLRPFERDISNVPFSPDGKPCTPPALNCYWPLNDYGFYTTTGIG

YQPYRVVVLSFELLNAPATVCGPKLSTDLIKNQCVNFNFNGLTGTGVLTPSSKRFQPFQQFGRDVSDFTD

SVRDPKTSEILDISPCSFGGVSVITPGTNASSEVAVLYQDVNCTDVSTAIHADQLTPAWRIYSTGNNVFQ

TQAGCLIGAEHVDTSYECDIPIGAGICASYHTVSLLRSTSQKSIVAYTMSLGADSSIAYSNNTIAIPTNF

SISITTEVMPVSMAKTSVDCNMYICGDSTECANLLLQYGSFCTQLNRALSGIAAEQDRNTREVFAQVKQM

YKTPTLKYFGGFNFSQILPDPLKPTKRSFIEDLLFNKVTLADAGFMKQYGECLGDINARDLICAQKFNGL

TVLPPLLTDDMIAAYTAALVSGTATAGWTFGAGAALQIPFAMQMAYRFNGIGVTQNVLYENQKQIANQFN

KAISQIQESLTTTSTALGKLQDVVNQNAQALNTLVKQLSSNFGAISSVLNDILSRLDKVEAEVQIDRLIT

GRLQSLQTYVTQQLIRAAEIMASANLAATKMSECVLGQSKRVDFCGKGYHLMSFPQAAPHGVVFLHVTYV

PSQERNFTTAPAICHEGKAYFPREGVFVFNGTSWFITQRNFFSPQIITTDNTFVSGNCDVVIGIINNTVY

DPLQPELDSFKEELDKYFKNHTSPDVDLGDISGINASVVNIQKEIDRLNEVAKNLNESLIDLQELGKYEQ

YIKWPWYVWLGFIAGLIAIVMVTILLCCMTSCCSCLKGACSCGSCCKFDEDDSEPVLKGVKLHYT

Alignment details:

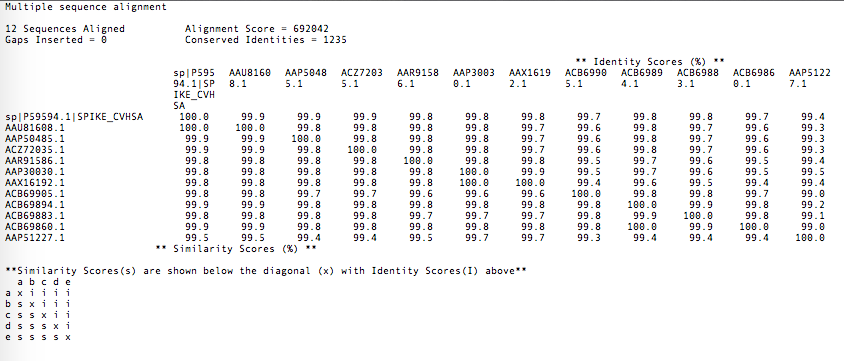

Sequence breakdown:

| **Accession #** | **Description** |
| --- | --- |
| P59594.1 | Spike glycoprotein |
| ACZ72035.1 | spike glycoprotein precursor [SARS coronavirus MA15] |
| AAP50485.1 | spike glycoprotein [SARS coronavirus FRA] |
| AAP30030.1 | spike glycoprotein S [SARS coronavirus BJ01] |
| AAP51227.1 | spike glycoprotein S [SARS coronavirus GD01] |
| AAX16192.1 | spike glycoprotein S [SARS coronavirus WH20] |
| ACB69905.1 | spike glycoprotein S [SARS coronavirus BJ182-12] |
| ACB69894.1 | spike glycoprotein S [SARS coronavirus BJ182-8] |
| ACB69883.1 | spike glycoprotein S [SARS coronavirus BJ182-4] |
| ACB69860.1 | spike glycoprotein S [SARS coronavirus BJ182a] |
| AAU81608.1 | S protein [SARS Coronavirus CDC#200301157] |
| AAR91586.1 | spike glycoprotein S [SARS coronavirus NS-1] |

Alignment: I will call specific attention to variable regions

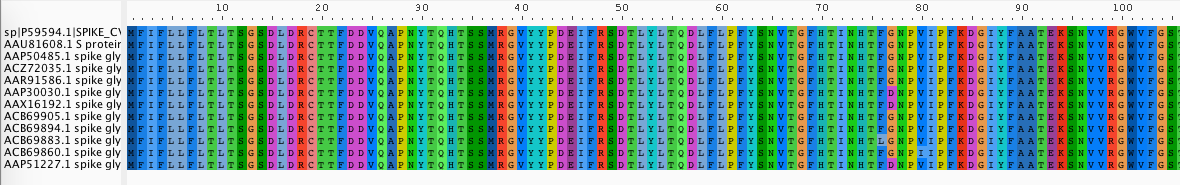

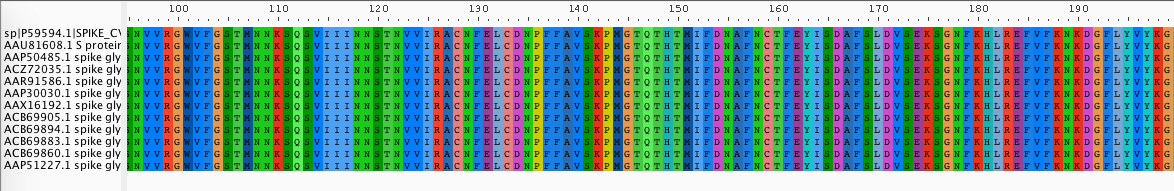

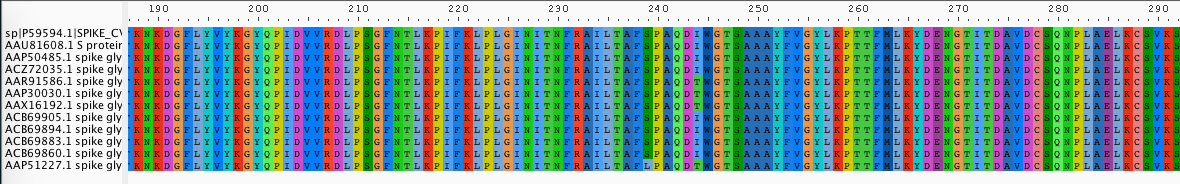

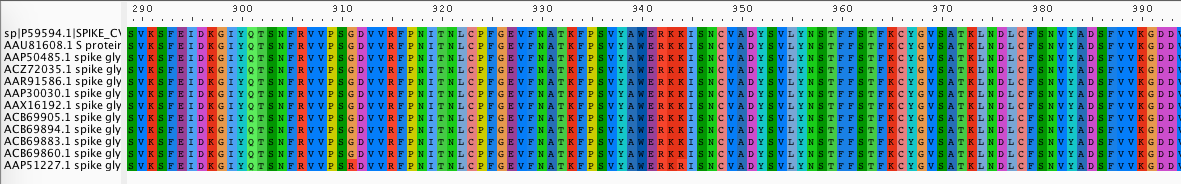

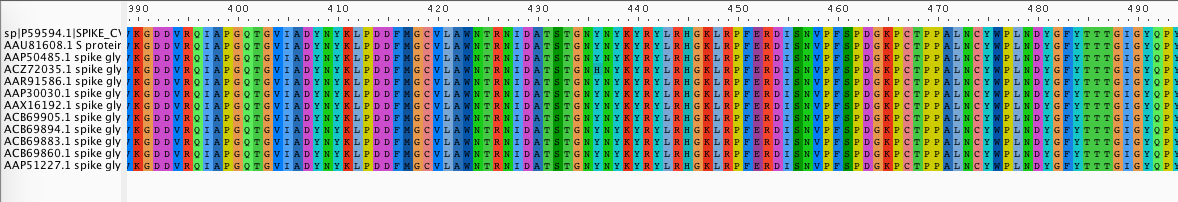

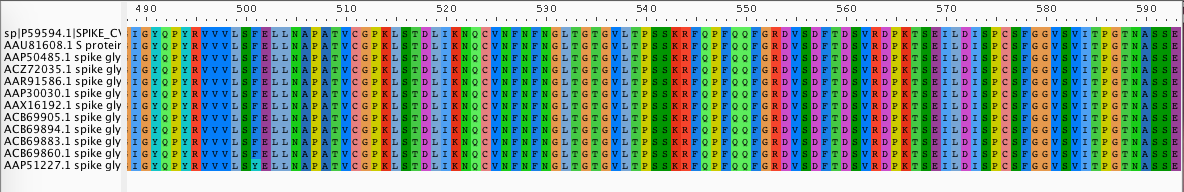

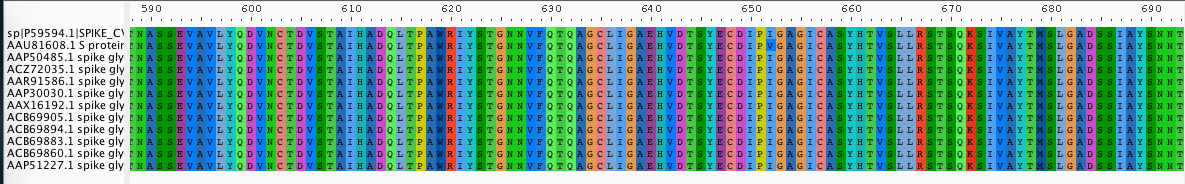

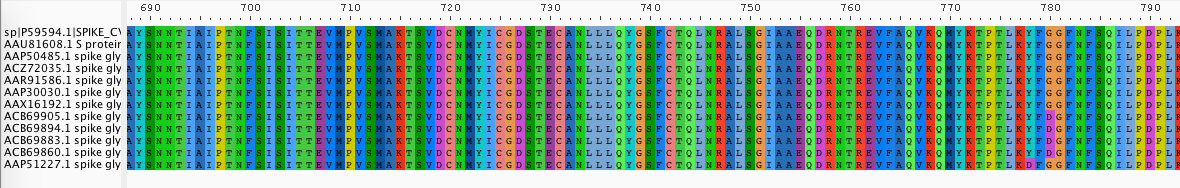

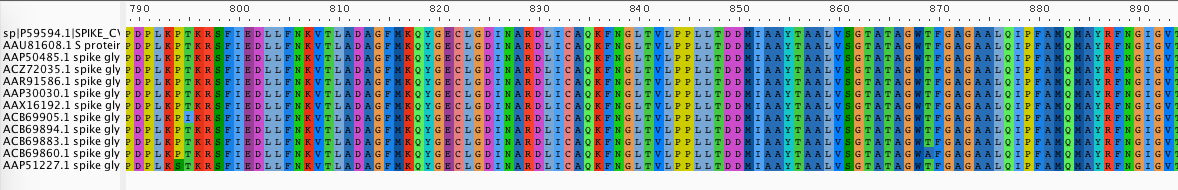

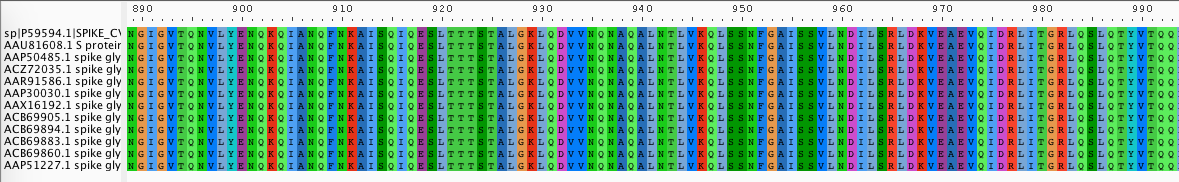

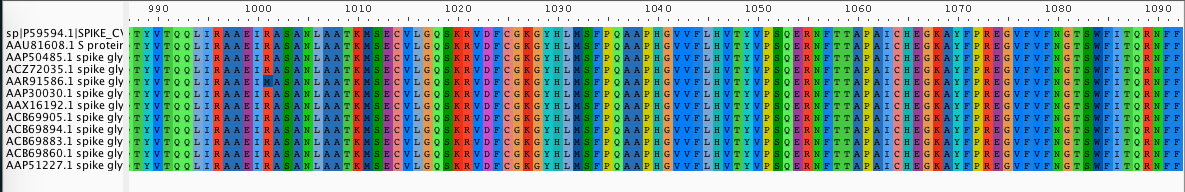

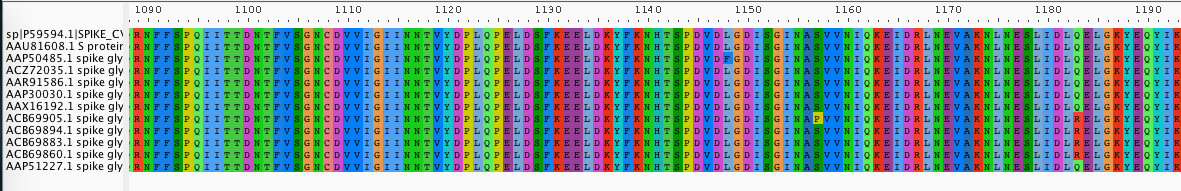

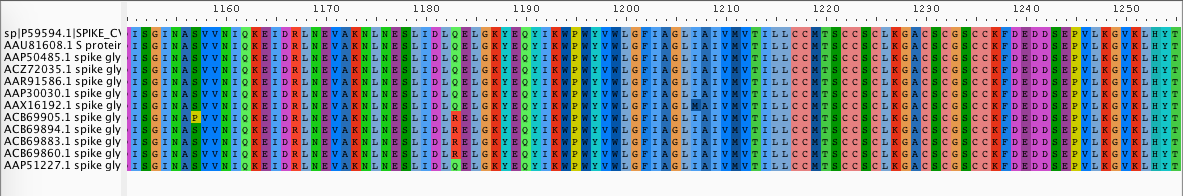
